# Histone H3 deacetylation promotes host cell viability for efficient infection by *Listeria monocytogenes*

**DOI:** 10.1101/2021.07.16.452630

**Authors:** Matthew J.G. Eldridge, Mélanie A. Hamon

## Abstract

For many intracellular bacterial pathogens manipulating host cell survival is essential for maintaining a replicative niche, and is a common strategy used to promote infection. The bacterial pathogen *Listeria monocytogenes* is well known to hijack host machinery for its own benefit, such as targeting the host histone H3 for modification by SIRT2. However, in what way this modification benefits infection, as well as the molecular players involved, remain unknown. Here we show that SIRT2 activity supports *Listeria* intracellular survival by maintaining genome integrity and host cell viability. This protective effect is dependent on H3K18 deacetylation, which safeguards the host genome by counteracting infection-induced DNA damage. Mechanistically, infection causes SIRT2 to interact with the nucleic acid binding protein TDP-43 and localise to genomic R-loops, where H3K18 deacetylation occurs. This work highlights novel functions of TDP-43 and R-loops during bacterial infection and identifies the mechanism through which *L. monocytogenes* co-opts SIRT2 to allow efficient infection.

## INTRODUCTION

The Sirtuin family (SIRT1-7) of NAD^+^-dependent deacetylases play key roles in many biological processes which are required to maintain cellular homeostasis, such as cell cycle, metabolism and DNA repair (Houtkooper *et al*, 2012; Gomes *et al*, 2015). Sirtuins have distinct subcellular localisations and divergent functional roles; however, they all regulate DNA and chromatin to various extents, particularly in response to DNA damage (Houtkooper *et al*, 2012). Despite their broad roles across different cellular compartments, mouse knockout models for all Sirtuins have emphasised their essential role in maintaining genome stability and cell survival in response to stress (Bosch-Presegué & Vaquero, 2014). As such, loss of individual Sirtuins is strongly associated with increased genome instability, and knockout mice often develop chromosomal aberrations and are predisposed to spontaneous tumorigenesis. The safeguarding of genomic stability by Sirtuins occurs in numerous ways including regulation of metabolic responses to stress, control of cell cycle checkpoints or adjustment of DNA damage signalling and repair through histone deacetylation (Bosch-Presegué & Vaquero, 2014).

Sirtuins 1, 6 and 7 display predominantly nuclear localisations, and as such have the most clearly defined roles in DNA damage responses. *Sirt1*^*-/-*^ cells have a reduced capacity to form DNA repair foci and fail to efficiently repair γ-radiation-induced DNA damage (Wang *et al*, 2008). This effect is believed to be driven by deregulation of chromatin dynamics via histone deacetylation, and repair proteins such as KU70 (Jeong *et al*, 2007), WRN (Chen *et al*, 2003) and XPA (Fan & Luo, 2010). SIRT6 can act as a DNA damage sensor which directly binds DNA breaks and promotes repair protein recruitment (Onn *et al*, 2020). Additionally, SIRT6 has been described to maintain the integrity of pericentric genomic regions through H3 lysine 18 (H3K18) deacetylation (Tasselli *et al*, 2016). Similarly, SIRT7 promotes the recruitment of the repair protein 53BP1 to sites of DNA damage which requires H3K18 deacetylation, and enhances non-homologous end joining (NHEJ) (Vazquez *et al*, 2016). However, SIRT7 lacks an ability to directly bind damaged DNA and instead requires Poly [ADP-ribose] polymerase 1 (PARP1) to localise to double strand breaks (Onn *et al*, 2020; Vazquez *et al*, 2016). By comparison, the mitochondrial Sirtuins have a more indirect role in preserving DNA stability. SIRT3 protects mtDNA by limiting mitochondrial superoxide levels (Kim *et al*, 2010) and positively regulating the DNA repair protein OGG1 (Cheng *et al*, 2013), while SIRT4 represses mitochondrial glutamine metabolism in response to genotoxic stress, thus promoting cell cycle arrest and allowing for more efficient DNA repair (Jeong *et al*, 2013).

Sirtuin 2 (SIRT2) is unique, as it is the only member of the family to hold a predominantly cytoplasmic localisation and have clear regulatory roles across multiple subcellular compartments, functioning in metabolism, cell cycle, inflammation, and oxidative stress responses (Lemos *et al*, 2017; Gomes *et al*, 2015; de Oliveira *et al*, 2012). However, SIRT2 is continuously shuttled between the cytosol and nuclear compartment, where it regulates nuclear proteins, such as p300 and p53 (Tanno *et al*, 2007; North & Verdin, 2007; Eldridge *et al*, 2020b; Peck *et al*, 2010; Black *et al*, 2008), and histones by deacetylation (Vaquero *et al*, 2006; Eskandarian *et al*, 2013). Furthermore, during mitosis SIRT2 accumulates in the nucleus, and becomes enriched at chromatin, where it deacetylates histone H4 lysine 16 (Inoue *et al*, 2007; Vaquero *et al*, 2006). As such, most of the described functions of SIRT2 in regulating DNA damage occur in the context of cell cycle progression and cell division. For instance, SIRT2-dependent H4K16 deacetylation has been shown to regulate H4K20me1 deposition, which in turn affects cell cycle checkpoint progression and reduces DNA damage accumulation during mitosis (Serrano *et al*, 2013). Similarly, SIRT2 promotes the activity of the anaphase-promoting complex/cyclosome (APC/C), which protects against mitotic catastrophe and promotes genome stability (Kim *et al*, 2011). Additionally, during the G2/M cell cycle checkpoint SIRT2 promotes CDK9 function which prevents the breakdown of stalled replication forks and arrests the cell cycle to allow additional time for DNA repair (Zhang *et al*, 2013). These reports point to SIRT2 having essential roles in maintaining genome stability which are linked to its nuclear accumulation during mitosis, but similar roles during interphase have not been shown. Our previous work identified a novel function of SIRT2 during infection with the bacterial pathogen *Listeria monocytogenes*. Infection triggers nuclear accumulation of SIRT2, where it becomes enriched on chromatin at transcriptional start sites (TSSs) of specific genes and induces deacetylation of H3K18 independently of the cell cycle (Eskandarian *et al*, 2013). Nuclear import of SIRT2 during infection is mediated in part by importin IPO7, and chromatin binding requires the dephosphorylation of SIRT2 at serine 25, allowing for H3K18 deacetylation (Pereira *et al*, 2018; Eldridge *et al*, 2020b). Importantly, SIRT2 activity at chromatin is essential for efficient *L. monocytogenes* infection in vitro and in vivo. However, how bacterial hijacking of SIRT2 promotes infection remains unknown. Given the roles of Sirtuins and H3K18 deacetylation in maintaining genome integrity, we reasoned that SIRT2 might function similarly during *L. monocytogenes* infection, in turn promoting host cell viability in order to better maintain the replicative niche (Ashida *et al*, 2011; Friedrich *et al*, 2017; Pirbhai *et al*, 2006; Behar & Briken, 2019; Knodler *et al*, 2005; Yan *et al*, 2009).

In this study we show that SIRT2 activity protects host cells from DNA damage and promotes host cell survival. We further show that the interaction with the DNA/RNA binding protein TDP-43 is essential for SIRT2 enrichment at the transcription start site (TSS) of specific genes and H3K18 deacetylation during infection. Mechanistically, we find that SIRT2 and TDP-43 function with DNA:RNA hybrids called R-loops to reduce the accumulation of host DNA damage caused by infection. Therefore, we show that during infection, the activity of SIRT2 on H3K18 is key in regulating cellular health, which is exploited by *L. monocytogenes* to maintain host genome integrity and cell viability thereby promoting infection.

## RESULTS

### SIRT2 activity maintains host cell viability during infection

Sirtuins have long been established to promote cell viability by maintaining genome stability. We were therefore interested in measuring cell viability during infection upon inhibition of SIRT2 activity. We performed an Alamar blue assay to measure the metabolic activity of HeLa cells, under uninfected and infected conditions, with and without SIRT2 inhibitor AGK2. Interestingly, infection with *L. monocytogenes* caused no reduction in host cell viability at either 6 or 24 hours post infection. However, in the presence of AGK2, a SIRT2 inhibitor, infected cells exhibited a significant reduction in viability (Fig. 1A). After 6 hours of infection, a slight 10% decrease in viability is detected in AGK2 treated cells, and by 24 hours cell viability is significantly decreased by 30% as compared with uninfected cells (Fig. 1A). Importantly, AGK2 treatment alone did not lead to a decrease in cell viability (Fig. 1A). Supporting this data, we performed cell counting assays at 6 h and 24 h post infection and were able to show that a higher proportion of dead cells were recovered at these time points (Fig. S1A). Therefore, although *L. monocytogenes* infection alone does not significantly impact cell viability, blocking SIRT2 activity during infection leads to significant cell death.

**Figure 1:**
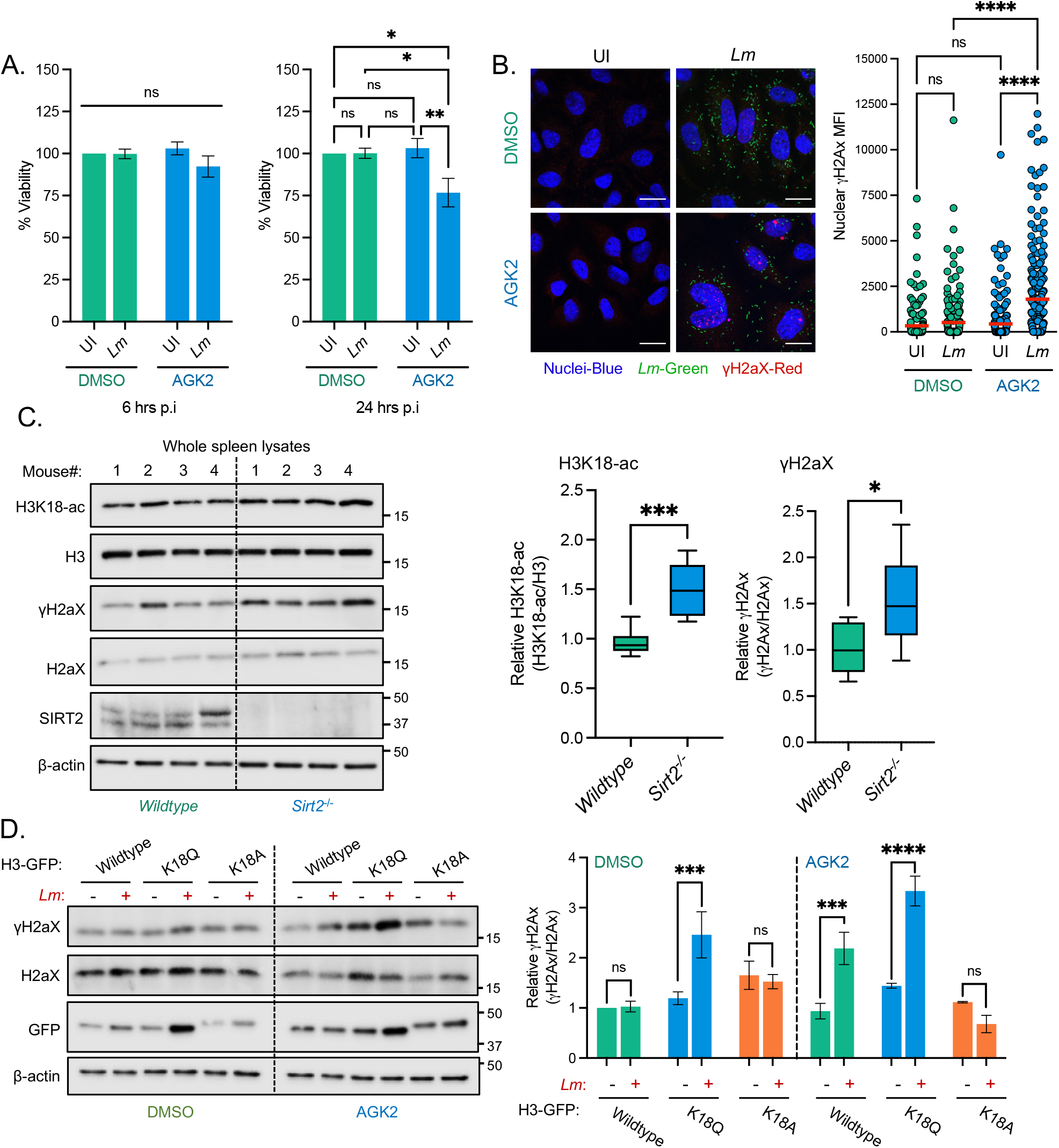
SIRT2 activity maintains host cell viability and genome integrity during *L. monocytogenes* infection. HeLa cells pre-treated for 2 hours in DMSO or 5 mM AGK2 were left uninfected (UI) or infected (EGD) with *L. monocytogenes* for 6 and 24 hours (A) or 24 hours (B). **(A)** Cytotoxicity was measured Alamar blue assay. Results are expressed as percent viability of uninfected cells. Plot shows mean ± SEM from three independent experiments. Statistical significance was determined by a Kruskal-Wallis test (ns = not significant, * = *p* < 0.05, ** = *p* < 0.01). **(B)** Representative images of immunofluorescence (left) detection of endogenous γH2aX (red) in HeLa cells left uninfected (UI) or infected for 24 hours with GFP-expressing *L. monocytogenes* (*Lm-* GFP). Scale bar is 20 µm. Quantification of nuclear γH2aX (right) from HeLa cells, data points represent the mean fluorescence intensity (MFI) of γH2aX within individual nuclei. Graphs display quantified nuclei from 2 independent experiments with the mean values of each condition represent by lines (red). Statistical significance was determined by one-way ANOVA with FDR Benjamini-Hochberg (BH) correction for multiple comparisons (ns = not significant, **** = *p* < <0.0001). **(C)** Immunoblot detection of stated proteins from infected mouse spleen lysates (left). Quantification of normalised H3K18-ac and γH2aX levels. Graphs show collated values from 8 mice from two independent experiments, box and whisker plot with solid line denoting the median value. Statistical significance was determined by Two-tailed Unpaired t test (* = *p* < 0.05, *** =*p* <0.001). **(D)** Immunoblot detection of γH2aX and total H2aX (left) from whole cell lysates of HeLa cells left uninfected (-) or infected with *L. monocytogenes* (EGD) for 24 hours. Cells are expressing stated H3-GFP plasmids and treated with DMSO or 5 mM AGK2. Images are representative of three independent experiments. Quantification of γH2aX levels (right) relative to uninfected. Results are expressed as intensity of actin normalised γH2aX bands relative to actin normalised total H2aX. Graph shows the mean ± SEM from three independent experiments. Statistical significance was determined by one-way ANOVA with Fisher’s LSD test (ns = not significant, *** =*p* <0.001, **** = *p* <0.0001).

### SIRT2 activity on H3K18 protects cells from infection-induced DNA damage

Since SIRT2 displays a cell protective effect during *L. monocytogenes* infection, we examined the consequences of SIRT2 inhibition on the DNA damage response. We monitored the accumulation of DNA damage by measuring the nuclear fluorescence intensity of the DNA damage marker γH2AX during late infection, in the presence or absence of the SIRT2 inhibitor AGK2.

Consistent with previous reports, *L. monocytogenes* infection induces low levels of DNA damage illustrated by an increase of γH2AX in host cell nuclei (Fig. 1B and S1B). At 24 hours post infection we observed a 15% increase in number of γH2AX positive cells as compared with uninfected conditions, accompanied by ∼1.5-fold increase in γH2AX mean fluorescence intensity (MFI) across the cell population (Fig. 1B, S1B and S1C). In uninfected cells treated with AGK2 there was no significant increase in γH2AX staining, suggesting that under resting conditions SIRT2 has no significant effect on the induction of DNA damage. By contrast, infected AGK2-treated cells accumulated significantly higher levels of DNA damage by 24 hours post infection, as evidenced by a 35% increase in the number of γH2AX positive cells and a concurrent ∼4-fold increase in the average nuclear γH2AX MFI (Fig. 1B, S1C and S1D). These data indicate that SIRT2 activity suppresses the accumulation of DNA damage during infection.

The impact of SIRT2 on infection-induced DNA damage was also determined in vivo. Spleens from wildtype and *Sirt2*^-/-^ mice were collected 72 hours after intravenous infection with *L. monocytogenes* and levels of γH2AX were assessed by immunoblotting. As expected, spleens from infected *Sirt2*^-/-^ mice had significantly higher levels of H3K18-ac and showed a trend towards lower bacterial numbers compared with wildtype mice (Fig. 1C and S1E). Similarly to what is observed during in vitro infection, levels of γH2AX were also significantly higher in *Sirt2*^-/-^ mice (Fig. 1C). Therefore, the role of SIRT2 in reducing DNA damage is detected in vivo, within organs that are targeted during infection.

We further wanted to determine whether it was the general activity of SIRT2 that was supressing DNA damage or the specific deacetylation of H3K18. To answer this question, we infected cells overexpressing GFP-tagged wildtype histone H3, or mutants where K18 was substituted with either glutamine (K18Q) or alanine (K18A) which respectively mimic acetylated and deacetylated H3K18. Under these conditions, DNA damage was measured by γH2AX immunoblotting. Upon transfection and expression of wildtype H3, DNA damage is observed only in infected cells that are AGK2-treated (Fig. 1D), similarly to what is observed by immunofluorescence under untransfected conditions (Fig. 1B). Alone, the expression of either mutant H3 K18Q or H3 K18A did not induce any significant increase in γH2AX levels in resting cells. Strikingly though, upon infection, expression of H3 K18Q is sufficient to induce higher levels of γH2AX (Fig. 1D), similar to the levels induced by AGK2 treatment. By contrast, expression of H3K18A does not increase γH2AX upon infection and, in fact, blocks γH2AX accumulation observed in AGK2 treated cells. These results suggest that deacetylation of H3K18 has a direct protective role against the accumulation of excessive DNA damage. Therefore, early recruitment of SIRT2 to DNA and its activity towards H3K18 is required to respond to infection-induced genotoxic stress.

#### SIRT2 interacts with TDP-43 for recruitment to chromatin

Our previous work showed that H3K18 deacetylation by SIRT2 occurs specifically at the TSSs of a subset of genes which are repressed during infection. However, SIRT2 does not display DNA binding properties. To identify interacting partners which could anchor SIRT2 to DNA we mined the previously published SIRT2 interactome (Eldridge *et al*, 2020b). Using the GeneCards database, we compiled lists of proteins known to interact with the TSSs of 5 different genes that are regulated by SIRT2 during infection (*MYLIP, ERCC5, LEF1, SYDE2, EHHADH*). We then compared these against the SIRT2 interactome to identify common proteins. One SIRT2-putative interactor which was common across all lists was TDP-43 (encoded by *TARDBP* gene) a DNA/RNA binding protein (Fig. S2). Further in silico analysis of previously identified infection-dependent SIRT2-repressed genes (Eskandarian *et al*, 2013) showed that 72% of these have TDP-43 present at their TSSs by ChIP-seq (ENCODE portal). Therefore TDP-43 represented a suitable candidate protein to recruit SIRT2 to chromatin at specific loci during *L. monocytogenes* infection.

To determine whether TDP-43 interacts with SIRT2 upon infection, HeLa cells were transfected with plasmids encoding GFP alone or GFP-tagged SIRT2 (SIRT2-GFP), then left uninfected or infected with *L. monocytogenes* followed by immunoprecipitation from isolated nuclei. Immunoblotting analysis showed that endogenous TDP-43 co-precipitates with SIRT2-GFP but not GFP alone in uninfected cells (Fig. 2A). Interestingly, following infection, TDP-43 binding to SIRT2 is further enriched by approximately 2-fold (Fig. 2A). Consistent with our previous interactome analysis these data show that a basal interaction between SIRT2 and TDP-43 occurs in the nuclei of uninfected cells, and we now show that this interaction is significantly enhanced in response to *L. monocytogenes* infection.

**Figure 2:**
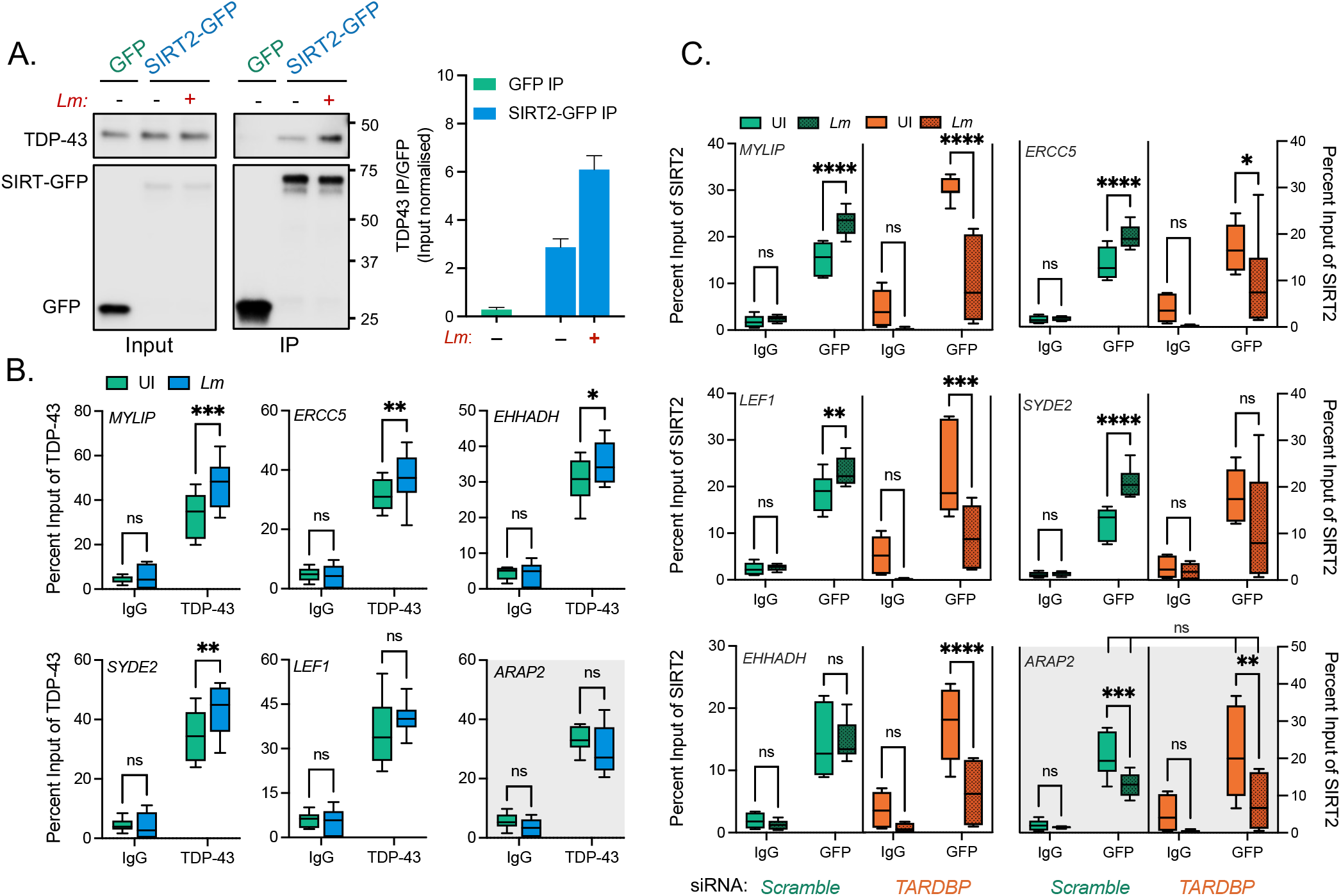
TDP-43 interacts with SIRT2 and is required for chromatin interactions and gene targeting upon infection. **(A)** HeLa cells expressing either GFP alone or SIRT2-GFP were left uninfected (-) or infected (+) for 3 hours. Cells were lysed and underwent immunoprecipitation using GFP-Trap® agarose beads. Cell lysates (Input) and IP fractions were immunoblotted using antibodies against GFP or TDP-43 (left). Quantification of endogenous TDP-43 enriched by GFP or SIRT2-GFP (right). Graph shows input normalised intensities of TDP-43 protein relative to SIRT2-GFP intensity detected from the same sample. Enrichment is expressed relative to basal interaction observed on uninfected cells. Graph shows the mean ± SEM from three independent experiments. **(B)** Chromatin immunoprecipitation (ChIP) using non-targeting control (Ctrl IgG) or TDP-43 (TDP-43 IgG) targeting antibodies quantified by qPCR. Chromatin was extracted from uninfected (UI - green) or infected (EGD - blue) HeLa cells 6 hours post infection. qPCR was carried out using primers targeting the transcriptional start sites of stated SIRT2-dependent or independent (gray background) genes. Graphs show collated technical readings (n=4) from three independent experiments and are presented as percent recovery of ChIP relative to input and plotted as box and whisker plot with solid line denoting the median value. Statistical significance determined by two-way ANOVA with FDR Benjamini-Hochberg (BH) correction for multiple comparisons (ns = not significant, * = *p* < 0.05, **= *p* < 0.01, *** =*p* <0.001). **(C)** Chromatin immunoprecipitation (ChIP) using non-targeting control (Ctrl IgG) or GFP (GFP IgG) targeting antibodies quantified by qPCR. Chromatin was extracted from HeLa cells stably expressing SIRT2-GFP and transfected with non-targeting *Scramble* (Green) or *TARDBP* targeting (Orange) siRNA. Cells were left uninfected (UI -clear) or infected (EGD - dotted) for 6 hours. qPCR was carried out using primers targeting the transcriptional start sites of stated SIRT2-dependent or independent (red box) genes. Graphs show collated technical readings (n=4) from three independent experiments and are presented as percent recovery of ChIP relative to input and plotted as box and whisker plot with solid line denoting the median value. Statistical significance determined by two-way ANOVA with FDR Benjamini-Hochberg (BH) correction for multiple comparisons (ns = not significant, * = *p* < 0.05, **= *p* < 0.01, *** =*p* <0.001, **** =*p* <0.0001).

We previously identified Ser25 as a residue on SIRT2 that was dephosphorylated upon infection and that this modification was necessary for SIRT2 to become enriched at chromatin (Pereira *et al*, 2018). Therefore, this post-translational modification could be involved in regulating the interaction between SIRT2 and TDP-43. To address this, we co-transfected HeLa cells with mCherry-TDP-43 and either WT SIRT2-GFP, phosphomimetic S25E SIRT2-GFP, or dephosphomimetic S25A SIRT2-GFP. Immunoprecipitation from cells with RFP-Trap beads followed by immunoblotting showed that all SIRT2 variants could interact TDP-43. Furthermore, both SIRT2 variants displayed an augmented interaction with TDP-43, particularly the S25A dephosphomimetic displayed a ∼2.5-fold increase in its interaction with TDP-43 as compared with the WT variant. This increase is similar to that observed following infection, suggesting that S25 dephosphorylation has a role in regulating this interface between SIRT2 and TDP-43 (Fig. S3).

We further wanted to establish whether TDP-43 was required for SIRT2-binding to DNA. Chromatin immunoprecipitation PCR (ChIP-PCR) of endogenous TDP-43 from uninfected HeLa cells showed that TDP-43 localises to the TSSs of SIRT2-regulated genes *MYLIP, ERRC5, LEF1, SYDE2* and *EHHADH*, consistent with multiple ChIP-seq data sets available from the ENCODE project database (Davis *et al*, 2018). Following *L. monocytogenes* infection TDP-43 shows a slight, and in most cases significant, enrichment of ∼10% at these genetic loci (Fig. 2B). By comparison *ARAP2*, a SIRT2 independent gene, does not show TDP-43 recruitment upon infection. To determine whether TDP-43 is necessary for the recruitment of SIRT2 to chromatin, we performed ChIP-PCR of GFP tagged SIRT2 from uninfected and infected cells transfected with either scramble or TDP-43 (*TARDBP*) targeting siRNA. For all tested genes, knockdown of TDP-43 does not change the basal level of SIRT2 recruitment in uninfected cells (Fig. 2C). As previously demonstrated, infection causes significant recruitment of SIRT2 to the TSSs of *MYLIP, ERRC5, LEF1, SYDE2* and *EHHADH* but not *ARAP2*. However, during infection, loss of TDP-43 from cells significantly reduces the recruitment of SIRT2 to the TSSs of these genes by ∼5-15%. By contrast, the SIRT2 activity-independent gene *ARAP2* shows a decrease in SIRT2 enrichment during infection which is not altered by the loss of TDP-43 (Fig. 2C). These data show that whilst TDP-43 is not required for the basal localisation of SIRT2 to chromatin in resting cells, the specific interaction and enrichment of SIRT2 which occurs following infection is dependent on TDP-43, and loss of TDP-43 dysregulates SIRT2-chromatin dynamics (Fig. 2C and S4B). Taken together these data show that infection enhances the interaction between SIRT2 and TDP-43 in the nucleus, and that TDP-43 is necessary for the infection-induced enrichment of SIRT2 at chromatin level to specific genetic locations.

### TDP-43 is required for SIRT2-dependent functions during infection

In the context of *L. monocytogenes* infection, our data strongly suggests that TDP-43 acts as a scaffold for SIRT2 recruitment to specific gene loci, and therefore would be essential for enabling SIRT2-dependent processes and related downstream phenotypes. 48 hours prior to infection HeLa cells were transfected with scrambled siRNA or a pool of three siRNAs which target either *SIRT2* or *TARDBP* mRNA, reducing their respective levels by ∼70% and 90% (Fig. S4A and S4B). As expected, in HeLa cells transfected with scramble siRNA, H3K18 deacetylation occurred normally during infection. Global H3K18-ac levels decreased by 30-40% as compared with uninfected cells (Fig. 3A). However, this decrease in acetylation levels was blocked in *TARDBP* silenced cells similarly to what was observed upon *SIRT2* silencing (Fig. 3A). Therefore, TDP-43 is required for SIRT2-dependent H3K18 deacetylation during infection.

**Figure 3:**
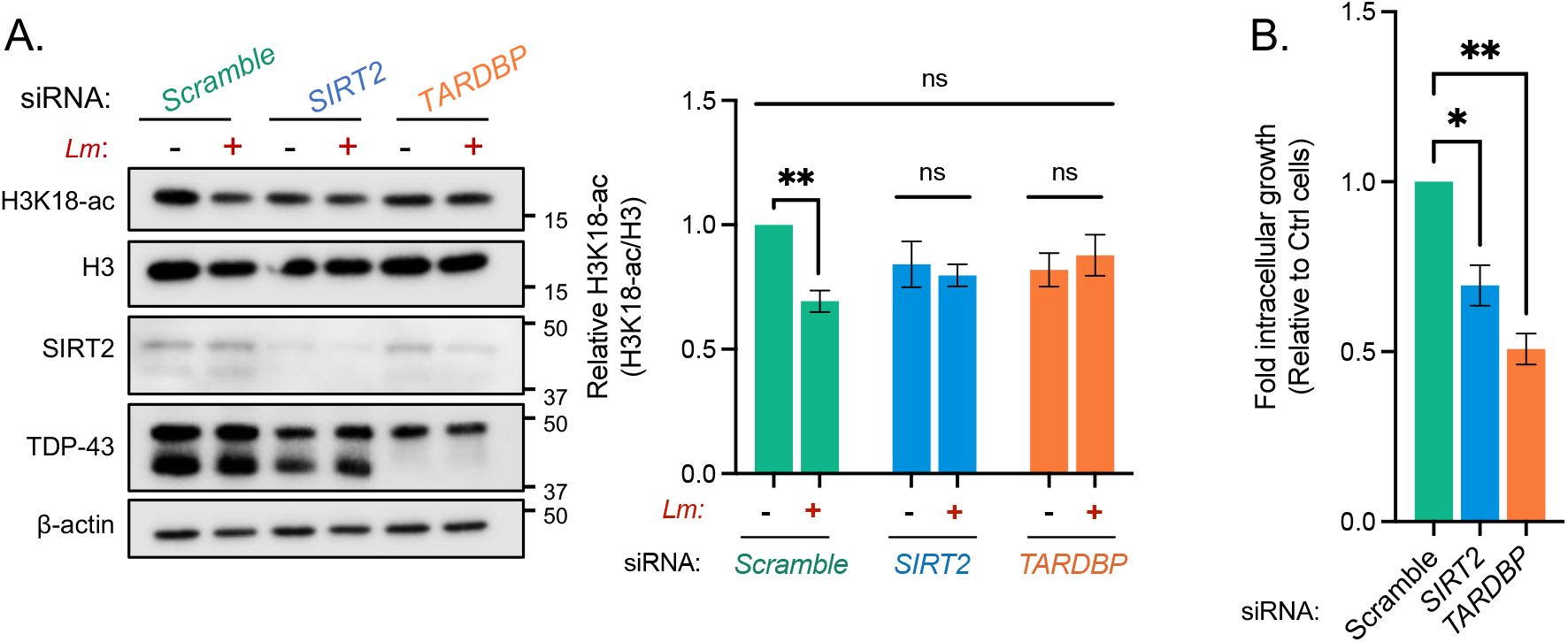
Silencing TDP-43 expression blocks H3K18 deacetylation and other SIRT2-related phenotypes during infection. **(A)** Representative image of H3K18 acetylation, SIRT2 and TDP-43 levels detected by immunoblotting (left) 6 hours post infection in uninfected HeLa (-) and *L. monocytogenes*–infected cells (+) transfected with stated siRNA. Quantification of H3K18 acetylation levels (right): band intensity of H3K18-ac and total H3 levels are normalised to β-actin followed by normalisation of H3K18-ac to total H3. Values are expressed as normalised band intensity relative to uninfected *Scramble* cells. Error bars represent SEM of four independent experiments. Statistical significance was determined by a Kruskal-Wallis test (ns = not significant, **= *p* < 0.01). **(B)** Fold change of intracellular *L. monocytogenes* colony forming units during infection of HeLa cells transfected with stated siRNAs. Data are presented as fold-change in recovered intracellular CFU between 2.5 and 24 hours post infection relative to *Scramble* siRNA cells. Graph shows the mean ± SEM from three independent experiments. Statistical significance was determined by a Kruskal-Wallis test (* = *p* < 0.05, **= *p* < 0.01).

We previously showed that SIRT2 activity is required to promote bacterial replication/survival in host cells, which is attenuated by enzymatic inhibition or genetic silencing of SIRT2 and results in lower recovered CFUs upon a 24h infection (Eskandarian *et al*, 2013; Pereira *et al*, 2018). Silencing of SIRT2 expression has no impact on early *Lm* invasion (Fig. S4C), however at later time points (24 hr p.i.) 30% fewer bacteria are recovered from cells transfected with SIRT2 siRNA compared with scramble controls (Fig. 3B) We performed similar experiments upon silencing of TDP-43 to assess *Lm* replication/survival during the later stages of infection in cultured cells. We obtained a similar reduction in bacterial numbers 24-hours post infection upon *TARDBP* knockdown as with *SIRT2* knockdown, where 50% fewer bacteria were recovered relative to scramble controls (Fig. 3B). Therefore, these results show that, like SIRT2, loss of TDP-43 has a negative impact on bacterial replication/survival within host cells and is therefore required to promote *Lm* infection. Altogether, our data demonstrate that TDP-43 is required for the execution of SIRT2-dependent H3K18 deacetylation during infection, and for the advantage SIRT2 can confer to *L. monocytogenes* during infection.

### R-loops are required for infection induced H3K18 deacetylation

TDP-43 is a nuclear DNA/RNA binding protein which specifically recognises single stranded nucleic acids. Recently, TDP-43 has been shown to interact with nucleic acid structures called R-loops, which preferentially form at TSSs when newly transcribed RNA anneals to the coding strand of DNA forming an RNA:DNA hybrid, and displaces a strand of ssDNA. To study the role of R-loops during infection, we overexpressed RNaseH1, an enzyme which resolves DNA/RNA-hybrids. Cells were transfected either with a mCherry control plasmid (pICE-mCherry-NLS) or a RNaseH1 expressing plasmid (pICE-RNaseH1-WT-NLS), and H3K18 deacetylation was monitored by immunoblotting. Expression of the control mCherry plasmid had no effect on the previously observed infection-induced H3K18 deacetylation. However, cells overexpressing RNaseH1 displayed no difference in acetylation levels, demonstrating that RNaseH1 expression blocks infection-induced deacetylation. In contrast, cells transfected with catalytically inactive mutant RNaseH1 ((pICE-RNaseH1-D10R, E48R-NLS) regained the ability to deacetylate H3K18 upon infection (Fig. 4A), which demonstrates that only catalytically active RNaseH1 blocks H3K18 deacetylation. These data therefore suggest that resolving of R-loops by expression of RNaseH1 blocks histone deacetylation, therefore indicating that the presence or formation of R-loops is required for this modification to occur.

**Figure 4:**
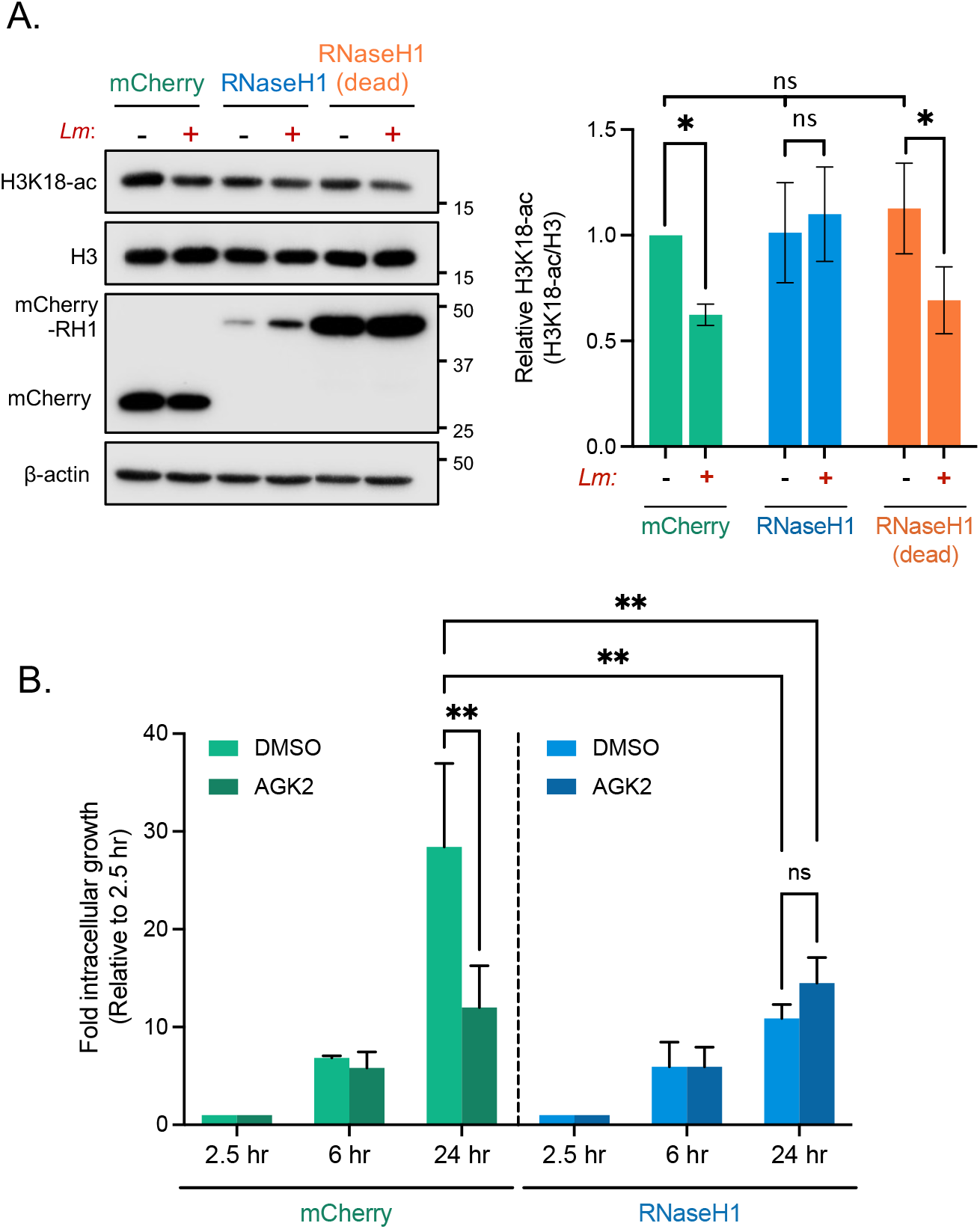
Blocking R-loop formation by overexpressing RNaseH1 inhibits infection induced H3K18 deacetylation and supresses bacterial intracellular survival. **(A)** Immunoblot analysis of H3K18 acetylation 6 hours post infection (left) in uninfected (-) and infected (+) HeLa cells expressing either mCherry, WT RNaseH1 or catalytically inactive RNaseH1 (dead). Quantification of H3K18 acetylation (right). H3K18-ac and total H3 levels band intensities are normalised to β-actin followed by normalisation of H3K18-ac to total H3. Values are expressed as normalised band intensity relative to uninfected mCherry cells. Error bars represent the SEM from at least three independent experiments. Statistical significance was determined by a Kruskal-Wallis test (ns = not significant, * = *p* < 0.05). **(B)** Fold change of intracellular *L. monocytogenes* colony forming units during infection of HeLa cells expressing stated plasmid constructs treated with wither DMSO or 5 mM AGK2. Data are presented as the fold-change in recovered intracellular CFU for each cell type at 6 and 24 hours post infection relative to their corresponding 2.5-hour timepoint. Graphs show the mean ± SEM from three independent experiments. Statistical significance was determined by two-way ANOVA with FDR Benjamini-Hochberg (BH) correction for multiple comparisons (ns = not significant, **= *p* < 0.01).

Similarly, we overexpressed RNaseH1 to determine whether resolving R-loops would influence the intracellular survival of *L. monocytogenes* as observed upon loss of SIRT2 or TDP-43 (Fig. 3B). In agreement with results from Figure 3B, cells transfected with a control mCherry plasmid behave as untransfected cells. By comparison, overexpression of RNaseH1 is alone sufficient to cause the same decrease in recovered bacterial colonies 24h post infection with no impact on infection at 6h (Fig. 4B and S5). Interestingly, additional SIRT2 inhibition with AGK2 does not have a cumulative effect on intracellular bacterial numbers, suggesting that R-loops are required for SIRT2 to promote infection (Fig. 4B). Together these data establish that R-loops are required for SIRT2 activity during infection, and that blocking their formation is alone sufficient to negatively affect the long-term survival of *L. monocytogenes* in host cells, phenotypically copying the loss of SIRT2 and TDP-43.

### TDP-43 and R-loops are required to protect against excessive infection induced DNA damage

Our data shows that SIRT2 activity and H3K18 deacetylation reduce the genotoxic effects of *L. monocytogenes* infection. We therefore asked whether TDP-43 and R-loops, which are also required for infection induced H3K18 deacetylation, would impact the accumulation of DNA damage. At earlier timepoints, where no infection-induced γH2AX is observed, loss of either SIRT2 or TDP-43 does not result in heightened γH2AX in cells (Fig. 5A). Consistent with our results using AGK2, infected cells depleted of SIRT2 or TDP-43 by RNAi display significantly elevated levels of γH2AX in infected cells, as detected by western blot at 24 hours post infection (Fig. 5B). Likewise, blocking the formation of R-loops by overexpressing RNaseH1 also significantly increases amount of γH2AX detected in infected cells at 24 hours post infection (Fig. 5C). This is in accordance with the role of R-loops in H3K18 deacetylation, demonstrating that R-loop inhibition phenotypically copies the loss of SIRT2 or TDP-43. Therefore, R-loops are required to protect infected cells from excessive DNA damage and are important for a productive *L. monocytogenes* infection.

**Figure 5:**
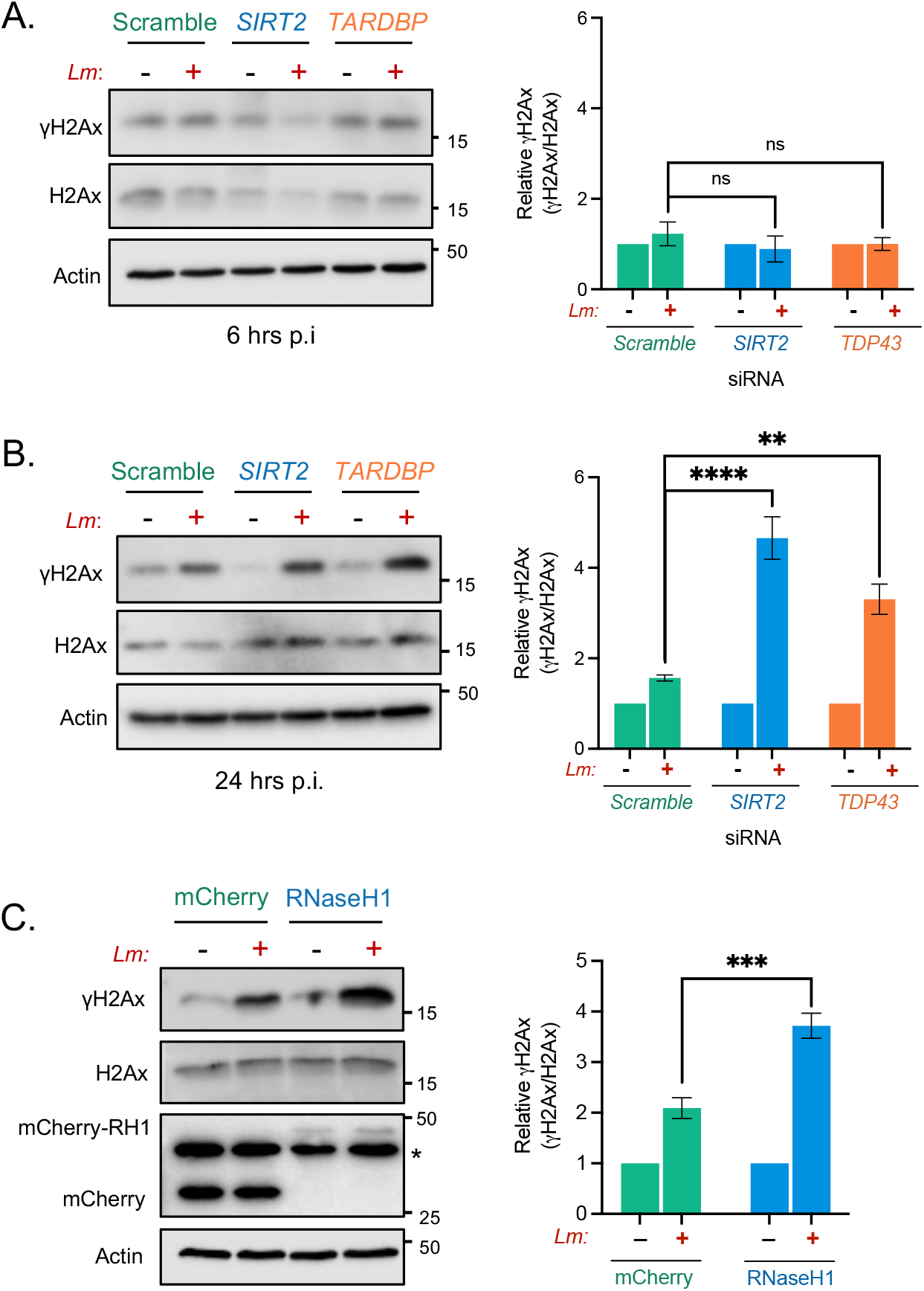
SIRT2, TDP-43 and R-loops function to protect cells from infection induced DNA damage. Immunoblot detection of γH2aX and total H2aX from whole cell lysates of HeLa cells transfected with stated siRNAs and left uninfected (-) or infected (+) for **(A)** 6 hours or **(B)** 24 hours. Quantified band intensities of γH2aX levels are present in graphs (right). Results are expressed as intensity of actin normalised γH2aX bands relative to actin normalised total H2aX. Graph shows the mean ± SEM from at least three independent experiments statis. Statistical significance was determined by one-way ANOVA with Dunnet correction for multiple comparisons (ns = not significant, ** =*p* <0.01, **** = *p* <0.0001). **(C)** HeLa cells expressing either mCherry or mCHerry-RNaseH1 were infected for 24 hours, immunoblot analyses and quantification of γH2aX (right) performed as stated for (A) and (B) *denotes non-specific band. Graph shows the mean ± SEM from five independent experiments. Statistical significance was determined by Two-tailed Unpaired t test (*** =*p* <0.001).

## DISCUSSION

Over the last decade, host nuclear factors and processes have been identified as common targets for bacterial pathogen manipulation during infection (Bierne & Hamon, 2020; Eldridge *et al*, 2020a; Dong & Hamon, 2020). Previous work demonstrated that, through InlB-induced signalling, *L. monocytogenes* triggers dephosphorylation of SIRT2 and co-opts its activity resulting in H3K18 deacetylation and augmented infection. In this study we decipher how SIRT2 interacts with chromatin upon infection and how its hijacking by *L. monocytogenes* contributes to bacterial infection. Here, we establish that SIRT2 activity towards H3K18-ac is required to maintain host cell health during infection, as in its absence host cell viability is reduced, resulting in a decrease in bacterial numbers. Specifically, SIRT2 activity and H3K18 deacetylation serve to protect host cell genome integrity by limiting the accumulation of DNA damage induced by *L. monocytogenes*. Mechanistically, infection and S25 dephosphorylation of SIRT2 enhance its interaction with the nucleic binding protein TDP-43 which enriches SIRT2 at the TSSs of specific genes to permit H3K18 deacetylation and maintain genome stability. These protective effects are also dependent on chromosomal DNA:RNA hybrids called R-loops which likely define the genomic locality of TDP-43 and thereby SIRT2 recruitment. Together, these data uncover a molecular mechanism involving a complex of SIRT2, TDP-43 and R-loops which regulate genomic integrity during infection and are the first to show functional roles for TDP-43 and R-loops in regulating cellular responses to bacteria (Fig. 6).

**Figure 6:**
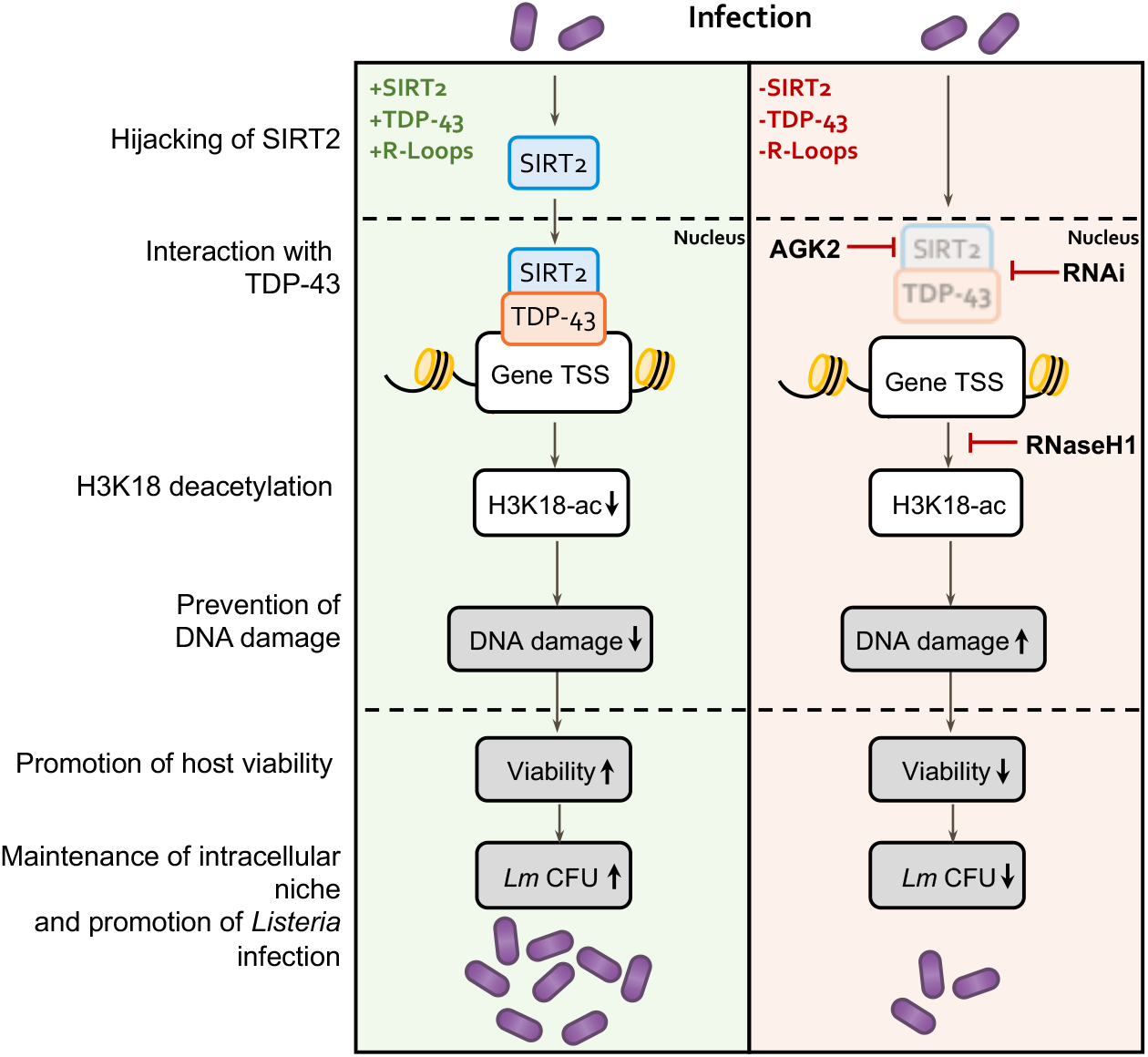
Model of host genome integrity protection by SIRT2. Schematic model of SIRT2 mechanisms of host cell protection and enhancement of infection. During infection, SIRT2 activity is hijacked by *Listeria* and is translocated to nucleus mediating H3K18 deacetylation. Recruitment to chromatin and histone deacetylation require TDP-43 and R-loops, which define the localisation of SIRT2 to specific genes. H3K18 deacetylation then directly functions to protect host genomic DNA from accumulating excessive DNA damage induced during infection by unknown mechanisms. This promotes genome integrity and cell viability thereby better supporting the intracellular lifestyle of *Listeria* and resulting in enhanced infection.

Our study defines H3K18 deacetylation by SIRT2 as the key factor required for protection from DNA damage during infection. Following exposure to ionizing radiation, SIRT7-mediated H3K18 deacetylation similarly protects genome stability by promoting the recruitment of the DNA repair protein 53BP1 and increasing the efficiency of NHEJ repair (Vazquez *et al*, 2016). Taken together, these data suggest that H3K18 deacetylation serves as a mark of DNA damage upon cellular stress for the recruitment of repair proteins. The general role of this mark in cellular stress needs to be investigated further.

Bacterial infection is a well-known inducer of DNA damage. However, some pathogens also target DNA damage responses in order to manipulate host cell fate, for instance to promote cell survival (Leitão *et al*, 2014; Samba-Louaka *et al*, 2014; Weitzman & Weitzman, 2014; Chumduri *et al*, 2013). *L. monocytogenes* infection has been reported to generate DNA damage in the host independently of ROS through an unknown mechanism (Samba-Louaka *et al*, 2014; Leitão *et al*, 2014). In fact, *L. monocytogenes* triggers degradation of the host DNA damage sensor MRE11, which promotes infection (Samba-Louaka *et al*, 2014). Previous work has suggested that this infection-induced DNA damage promotes infection by delaying host cell cycle progression and increasing the host cellular nucleotide pool which can be scavenged by bacteria, promoting their replication (Leitão *et al*, 2014). In macrophages, *L. monocytogenes* infection induces DNA breaks which are generated by nitric oxide production in responses to TLR signalling. This in turn activates a DNA damage response (DDR) pathway that regulates a pro-inflammatory transcriptional program to augment macrophage responses (Morales *et al*, 2017). Consistent with its potential role in DDR during infection, our previously published SIRT2 interactome identified many DNA damage sensor and repair proteins including KU70/ KU80, RPA1, FEN1 and PARP1 (Eldridge *et al*, 2020b).Irrespective of the role that DNA damage might play during infection, mitigating its cytotoxic effects would benefit the maintenance of the intracellular niche.

The exploitation of SIRT2 requires the effector InlB which binds and activates the host receptor for hepatocyte growth factor (HGF) c-Met (Eskandarian *et al*, 2013). Classically this interaction is recognised to induce the uptake of *L. monocytogenes* into non-phagocytic cells by clathrin-mediated endocytosis and trigger host pro-survival signalling through PI3K and AKT which are also typical of HGF stimulation to promote infection (Radoshevich & Cossart, 2018). In non-infectious pathologies such as cancer the HGF/c-Met axis is often hyperactive which greatly contributes to oncogenesis by promoting cancer cells survival. Additionally, constitutive c-Met signalling in cancer cells also stimulates multiple DNA repair mechanisms which can render tumours resistant to anti-cancer drugs which act by inducing DNA damage (Medová *et al*, 2014; Comoglio *et al*, 2018; De Bacco *et al*, 2016; Li *et al*, 2009). Interestingly, mutant *ΔinlB L. monocytogenes* induce higher levels of host DNA damage during infection despite being less invasive (Samba-Louaka *et al*, 2014). As such, the engagement of c-Met, hijacking of SIRT2, and subsequent H3K18 deacetylation, could represent a specific DDR mechanism which is exploited by *L. monocytogenes* to promote host cell survival. TDP-43 is a ubiquitously expressed protein belonging to the heterogenous nuclear ribonucleoprotein (hnRNP) family which has specificity for single stranded TG/UG-rich DNA and RNA (Kitamura *et al*, 2018; Kuo *et al*, 2014; Buratti *et al*, 2004; Buratti & Baralle, 2001). Like SIRT2, TDP-43 also shuttles between the cytoplasm and nucleus (Ayala *et al*, 2008), however, it primarily maintains a nuclear localisation and functions in RNA processing and as a direct transcriptional repressor (Lagier-Tourenne *et al*, 2010; Lalmansingh *et al*, 2011). Though primarily monomeric physiological oligomerisation of TDP-43 has also been described and is believed to regulate DNA-binding and stress resistance (Chang *et al*, 2012; Afroz *et al*, 2017). TDP-43 has been identified as a causative factor of the neurodegenerative disease amyotrophic lateral sclerosis (ALS), mostly commonly due to mutations which cause it to mislocalise to the cytoplasm and self-assemble into large prion-like aggregates (Jo *et al*, 2020). As well as their direct pathological roles, ALS-related mutations also disrupt the native functions of TDP-43 revealing that it acts as a scaffold for the recruitment of DNA repair proteins. As such, ALS mutant or TDP-43 deficient neuronal cells have defects in NHEJ DNA repair and are more sensitive to genotoxic agents (Mitra *et al*, 2019; Konopka *et al*, 2020). The interaction of TDP-43 with SIRT2 had not previously been shown, however our work suggests that SIRT2 could have an important role in regulating DNA repair in ALS

R-Loops also regulate DNA damage; however, whether they are detrimental or beneficial for the maintenance of genome integrity remains controversial (Marnef & Legube, 2021; Crossley et al, 2019; Niehrs & Luke, 2020). Persistent R-loops have been demonstrated to cause DNA damage due to incorrect processing by nucleotide excision repair nucleases XPG and XPA or by blocking replication fork progression resulting in the formation double strand breaks (Gan et al, 2011; Cristini et al, 2019). However, R-loops can also function to promote DNA repair, particularly in the context of transcriptionally coupled homologous recombination repair and NHEJ (Marnef & Legube, 2021; Chakraborty et al, 2016; Yasuhara et al, 2018). We find that blocking R-loop formation by overexpressing RNaseH1 leads to higher levels host DNA damage in response to *L. monocytogenes*, suggesting that R-loops play a protective role during infection.

Interestingly, in silico analysis shows that many SIRT2 regulated sequences contain or are predicted to contain R-loops; additionally there are multiple studies which demonstrate that TDP-43 localises to and interacts with R-loops (Gianini et al, 2020; Mosler et al, 2021; Cristini et al, 2018). Our data suggest that, in the context of *L. monocytogenes* infection, TDP-43 recruits SIRT2 to chromatin, as it can for other DDR factors (Mitra *et al*, 2019; Konopka *et al*, 2020). Consistent with this, inhibition of H3K18 deacetylation by RNaseH1 indicates that R-loops act upstream of SIRT2 activity, suggesting that R-loops are recognised by TDP-43, which serves as a platform for SIRT2 recruitment during infection.

Sirtuins have long been known to regulate cellular responses to DNA damage. Recent work showed that SIRT6 acts as a direct DNA damage sensor whose activity initiates DNA repair responses when localised to broken DNA. Interestingly, although SIRT2 lacks an ability to directly bind DNA, SIRT2 fused with a lactose repressor (LacR) element (to allow DNA binding) showed that recruitment of SIRT2 to DNA was sufficient to initiate the recruitment of DNA repair proteins (Onn *et al*, 2020). Given that SIRT2 has also been shown to promote mycobacterial infection, this interaction with TDP-43 could function during other bacterial infections (Bhaskar *et al*, 2020). Independently, these factors are also linked to many non-infectious human pathologies, and mutations in SIRT2, TDP-43 and R-loop regulating factors have been linked with age-related illnesses such as cancer and neurodegenerative diseases, both of which are also intrinsically linked to the deregulation of DNA damage responses. As such, these mechanisms not only have implications in better understanding cellular response to infection but could also extend to other factors of human health and disease.

## EXPERIMENTAL PROCEDURES

### Cell Culture, inhibitor treatments and Listeria monocytogenes infections

HeLa (ATCC, CCL-2) cells were grown to semi-confluency in minimum essential medium (MEM) plus GlutaMAX (Gibco) supplemented with 1 mM sodium pyruvate (Gibco), and 10% fetal bovine serum (FBS). 24 hours before infection, HeLa cell medium was changed to low serum (0.25% FBS) MEM medium containing 1 mM sodium pyruvate. *Listeria monocytogenes* EGD (see Supplementary Table S1) were grown overnight in brain heart infusion (BHI) liquid broth with shaking at 37°C. For infection bacteria were subcultured (1 in 10) into fresh BHI and grown to mid log phase (OD_600_ = 0.8-1) and washed 3× in MEM + 0.25% FBS before being added to cells. Bacteria were then added onto cells at a MOI of 100 (unless otherwise stated) and incubated for 1 hour. Cells were then washed 3× in MEM + 0.25% FBS and incubated in fresh medium for 30 minutes prior to the addition of 10 µg.mL^-1^ gentamicin for the remaining time of the infection. Inhibitors were added 2 hours prior to infection and remained present until 1 hour post infection when cells were washed. SIRT2 inhibitor AGK2 (Calbiochem) was used at a concentration of 5 mM.

### Cell viability alamar Blue assay

Cells were incubated at 37°C in fresh medium containing 10% alamarBlue reagent for 1-2 hours. Fluorescence (Ex/Em 560/590 nm) was then read using a Cytation 5 (BioTek). Fluorescence readings were blank corrected to wells containing only culture medium and results are expressed as a percentage of uninfected cells viability.

### Immunofluorescence microscopy

For immunofluorescence HeLa cells were plated onto coverslips prior to treatments. Following treatments cells were washed three times in PBS and fixed using 4% PFA in DPBD for 10 mins. Cells were then permeabilised for 10 mins in 0.2% Trition X-100 PBS. Coverlips were then incubated in blocking buffer (1% BSA TBS) for 1 hour. For immunostaining coverslips were inverted on to droplets of blocking buffer containing Phospho-Histone H2A.X (Ser139) (CST, 2577) antibody (1:500) then incubated in a humidified chamber overnight at 4°C. Subsequently, coverslips were washed 3 times in PBS + 0.1% Tween then incubated at room temperature in the dark for 1 hour in blocking buffer containing Alexa Fluor 546 goat anti-rabbit IgG (Invitrogen, A-11035) secondary antibody (1:1500) for 1 hour. Coverslips were washed three times TBS + 0.1% Tween, nuclei were stained with 300 nM (100 ng.mL^-1^) Hoechst 33342 for 15 mins. Coverslips were then washed three times in TBS, rinsed briefly in distilled water and mounted using Fluoromount-G® Mounting Medium (INTERCHIM). All images were acquired using a Zeiss Axio Observer spinning-disk confocal microscope equipped driven by the MetaMorph software. For quantification a minimum of ten fields of view were obtained per condition of each biological replicate.

### Immunoblotting and band quantification

Cell lysates were prepared in 2× Laemmli loading buffer supplemented with cOmplete protease inhibitor and PhosSTOP phosphatase inhibitor tablets (Roche), 1 mM PMSF, 5 mM sodium butyrate and 5% β -mercaptoethanol. Proteins were separated by SDS-PAGE using TrisGlycine buffer systems and transferred to PVDF membranes (Bio-Rad Laboratories). Membranes were blocked for 1 hour in TBS + 0.1% Tween containing 5% milk and then incubated with primary antibodies (as per the manufactures instructions) overnight at 4°C with rocking. Immunoblot quantification used images acquired on a Chemidoc MP (Bio-Rad), analyzed using Image Lab software (Bio-Rad Laboratories).

### Antibodies

Antibodies used in this study are as follow; anti-GFP antibody (Abcam, ab290), Acetyl-Histone H3 (Lys18) antibody (CST, 9675), anti-Histone H3 antibody (Abacam, ab1791), anti-β-actin (Sigma, AC-15), anti-TDP-43 antibody (Sigma, T1705), anti-SIRT2 (CST, 12650) anti-mCherry antibody (1C51) (Novus Biologicals, NBP1-96752), anti-γH2A.X (S139) antibody (2OE3) (CST, 9718S), anti-H2A.X antibody (CST, 2595S), Phospho-Histone H2A.X (Ser139) antibody (immunofluorescence) (CST, 2577).

### In vivo animal studies

Protocols for animal studies were reviewed and approved by the Comité d’Ethique pour l’Expérimentation Animale of Institut Pasteur under approval number Dap170005 and performed in accordance with national laws and institutional guidelines for animal care and use. Wild-type C57BL/6 mice were purchased from Janvier Labs. *Sirt2*^tm1a(EUCOMM)Wtsi^ mice were obtained from the Sanger Center. For details, see www.informatics.jax.org/javawi2/servlet/WIFetch?page=alleleDetail&key=606707. Female mice aged 8–16 weeks old were infected by intravenous injection of 10^5^ bacteria per animal and proceeded for 72 hours.

### RNA interference and DNA transfections

Transient RNAi was carried out using ON-TARGETplus siRNAs from Dharmacon. HeLa cells were transfected with siRNA targeting either *SIRT2* (SMARTpool L-004826-00-0005), or *TARDBP* (SMARTpool L-012394-00-0005). ON-TARGETplus Non-targeting Pool siRNA (D-001810-10-05) served as the negative control. Reverse transfections were performed in 6 well plates using Lipofectamine RNAiMAX reagent (Invitrogen). Briefly, 2.5×10^5^ HeLa were added to wells containing 15 pmol of siRNA mixed with 3 µL Lipofectamine RNAiMAX in 500 µL OptiMEM (Gibco) and incubated for 48 hours prior to further treatment or infection.

Transient expression of DNA plasmids was carried out in 6 well plates by reverse transfection using Lipofectamine LTX (Invitrogen). Briefly, 5-6×10^5^ HeLa cells were added to wells containing DNA-lipid complexes consisting of 1 µg plasmid DNA mixed with 1.5 µL Plus reagent and 3 µL LTX transfection reagent in 500 µL OptiMEM.

### RNaseH1 transfection and induction

For experiments testing the role RNaseH1 overexpression HeLa stably express the tetracycline repressor (HeLa T-Rex) protein (Agathe Subtil) were used to enable induction of pICE plasmids. HeLa T-Rex cells were transfected as described above. For plasmid induction transfected cells were incubated overnight with 10 ng.mL^-1^ Anhydrotetracycline hydrochloride (AHT).

### Co-immunoprecipitation with MNase lysis

Immunoprecipitations of SIRT2-GFP were performed using GFP-Trap® agarose beads (Chromotek). Briefly, 2-4×10^6^ HeLa cells were transfected with tagged-SIRT2 or empty pEGFP-N1/pmCherry-C1. 24 hours post transfection cells were collected using PBS+EDTA washed once in PBS and resuspended in 100uL MNase reaction buffer (1mM CaCl_2_, 0.2% NP-40, 50mM Tris-HCl (pH 7.6) 1mM CaCl_2_, 0.2% NP-40, 50mM Tris-HCl (pH 7.6) and 10 U micrococcal nuclease to react at 37 °Cfor 20 min. Reaction was terminated with 5 mM EDTA and sample was diluted 1:1 with 2X RIPA buffer containing 1 mM PMSF and sodium butyrate and incubated on ice for 10 minutes. Lysate was cleared by centrifugation at 20000 *xg* for 5 minutes and the resulting supernatant was diluted with 600 µL of wash/dilution buffer (10 mM Tris/Cl pH 7.5; 150 mM NaCl; 0.5 mM EDTA). 40 µL was removed for input and the remaining lysate was incubated with GFP-Trap® agarose beads at 4°C with agitation for 1 hour. The beads were washed twice in wash buffer and once in wash buffer containing 300 mM NaCl. Proteins were eluted by boiling beads in 50 µL 2× Laemmli buffer with 5% β-mercaptoethanol.

### Chromatin immunoprecipitation PCR

3-5⨯10^6^ cells were cross-linked at room temperature with 1% formaldehyde for 10 minutes followed by quenching with 130 mM glycine for 5 minutes. Chromatin extraction and ChIP-PCR were performed as previously described with slight modifications (Connor *et al*, 2021). Briefly, cell pellets were lysed on ice in nuclear isolation buffer (NIB) supplemented with 0.2% Triton X-100 and inhibitors (1× cOmplete™ protease, 1X PhosSTOP™, 10 mM sodium butyrate, 0.2mM PMSF) for 30 min with gentle pipetting every 10 min. Nuclei were collected by centrifugation and re-suspended in chromatin shearing buffer with inhibitors. Chromatin was fragmented by sonication (30 cycles of 15 s ‘on’ and 30 s ‘off’) with a Bioruptor (Diagenode) to 200 – 1000 bp. Sheared chromatin was cleared by centrifugation, sampled for size using 2% agarose gel electrophoresis and quantified using Pico488 (Lumiprobe, 42010). 2 µg of antibody (anti-TDP-43, T1705; anti-GFP antibody, ab290) was used per ChIP and were bound to Dynabeads Protein G (Invitrogen) overnight at 4°C with gentle rotation. Chromatin was diluted to 10-15 µg/IP with SDS dilution buffer supplemented with inhibitors. 8% of ChIP sample volume was reserved to serve as input. Diluted chromatin was then added to antibody bound Dynabeads and incubated at 4°C overnight with gentle rotation. IP samples were washed sequentially with 1 mL of buffers 1–6. Water containing 10% Chelex was added to washed beads and input samples and were eluted and de-crosslinked by boiling for 10 minutes. Samples were then treated with RNase A at room temperature for 10 minutes at 37°C followed by proteinase K (500 μg/ml) for 20 min at 55°C. Samples were then boiled for a further 10 min and recovered DNA was purified by phenol–chloroform extraction and isopropanol precipitation and resuspended in molecular grade water. ChIP DNA was quantified by qRT-PCR using iTaq™ Universal SYBR® Green Supermix and results were expressed as percent recovery from input calculated as 2 raised to cycle adjusted input sample quantitation cycle (Cq) value minus the Cq immunoprecipitation sample, multiplied by 100. For buffer formulations and primer sequences see Supplementary Tables S2 and S3 respectively.

### RNA isolation, reverse transcription, and qRT-PCR

RNA was extracted from cells using TRIzol™ Reagent (Life Technologies) extraction method as per the manufacturer’s instructions. cDNA was synthesised from 2 µg purified RNA using iScript™ cDNA Synthesis Kit (Bio-Rad) and quantified by qRT-PCR using iTaq™ Universal SYBR® Green Supermix. Data was analysed using ΔCT method relative to *GAPDH*.

### Plasmids Single, oligo mutagenesis and molecular cloning

Routine cloning was carried out by sequence- and ligation-independent cloning (SLIC) (Jeong *et al*, 2012) for primers see Supplementary Table S3. For further details on plasmids used in this study see Supplementary Table S4. pEGFP-H3 WT, pEGFP-H3 K18A, pEGFP-H3 K18Q were a gift from Dr Fang-Lin Sun (Liu *et al*, 2012). pmCherry TDP-43 was cloned from TDP43 NOTAG1(Addgene #28206) which was a gift from Zuoshang Xu (Yang *et al*, 2010). pICE-NLS-mCherry (Addgene #60364) and pICE-RNaseHI-WT-NLS-mCherry (Addgene #60365) were gifts from Patrick Calsou (Britton *et al*, 2014). pICE-RNaseHI-WT-NLS-mCherry (Dead) was made by introducing inactivating D10R and E48R mutations by single oligo mutagenesis (Shenoy & Visweswariah, 2003).

### Statistical analysis

All experiments were repeated at least twice, and statistical tests are reported in the figure legends. Data normality was tested by Shapiro-Wilk test, and appropriate parametric or non-parametric tests were used. Data plots and statistics were generated using Prism (version 9, GraphPad Software Inc.).

## ACKNOWLEDGEMENTS

We would like to thank and acknowledge Pascale Cossart for her support during the project and her contribution towards the acquisition of financial support. We also thank Michael G. Connor for his help processing samples for ChIP-PCR, Marie-Anne Nahori for performing mouse infections and Tiphaine Marie-Noelle Camarasa for help processing mouse organs. We thank Julia Sanchez-Garrido, Pascale Cossart, Julia Torne Cortada and Tiphaine Marie-Noelle for critical reading of the manuscript. MJGE is supported by a fellowship from the French Government’s Investissement d’Avenir program, the Laboratoire d’Excellence ‘‘Integrative Biology of Emerging Infectious Diseases’’ (ANR-10-LABX-62-IBEID). Work in the M.A.H. laboratory received financial support from the Institut Pasteur, the National Research Agency (ANR-EPIBACTIN), the Fondation pour la Recherche Médicale (FRM), the Fondation iXCore-iXLife and the Pasteur-Weizmann research fund.

## AUTHOR CONTRIBUTIONS

M.J.G.E., and M.A.H. conceived and designed the experiments. M.J.G.E. conducted the experiments. M.J.G.E. analysed results. M.J.G.E. wrote the original manuscript draft. M.J.G.E. and M.A.H. edited and reviewed the manuscript. M.A.H. supervised the research and obtained funding. All authors approved the final manuscript.

## FIGURE LEGENDS

**Figure S1:**
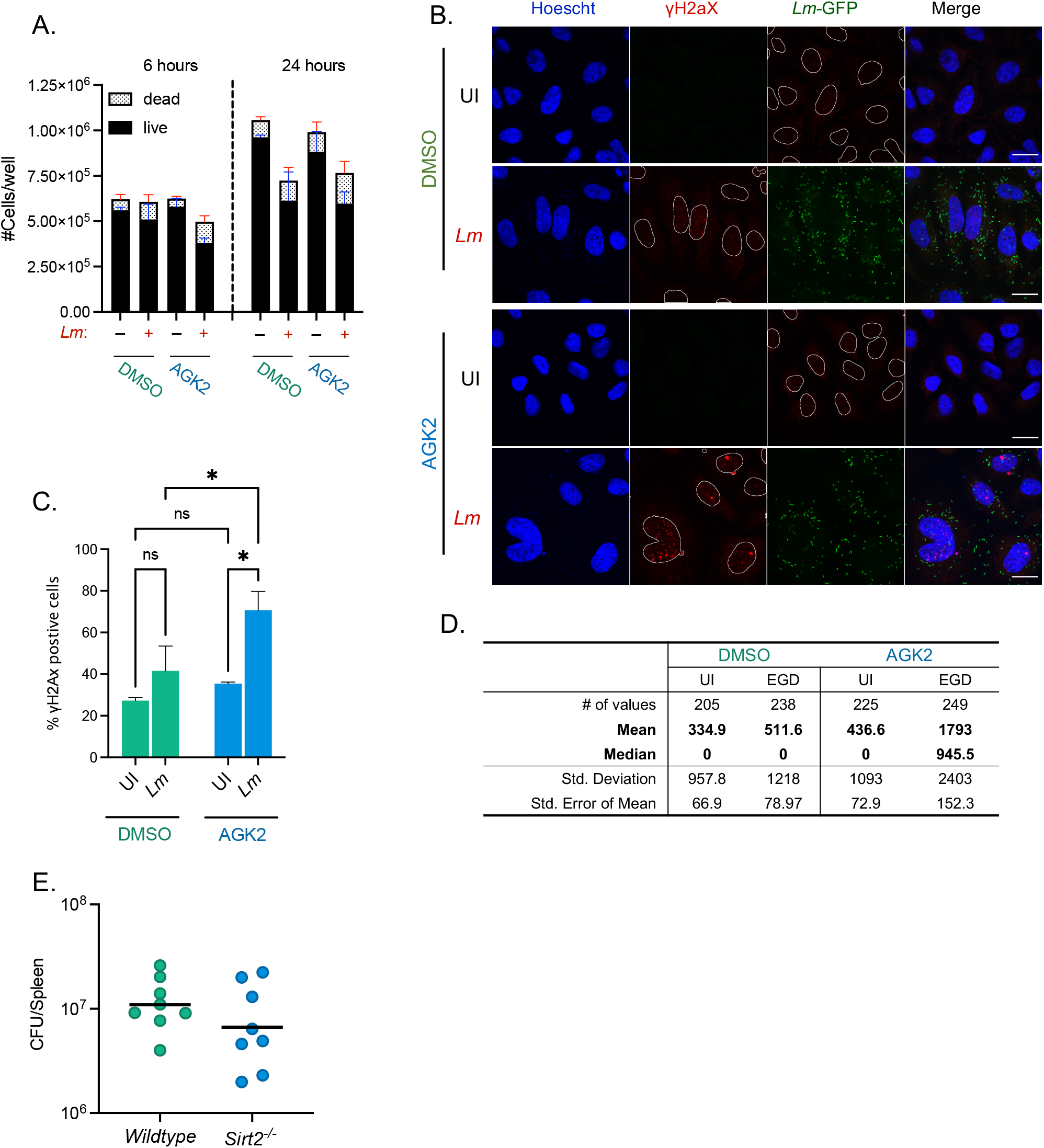
SIRT2 inhibition effects cell viability and DNA damage during infection. HeLa cells pre-treated for 2 hours with DMSO or 5 mM AGK2 (A-D). **(A)** Enumeration of live (Trypan negative) and dead (Trypan positive) cells at stated times post infection. Cells were enumerated with Countess™ II Automated Cell Counter from 2 independent experiments with the me. **(B)** Reprehensive unmerged images of nuclear γH2aX from uninfected and infected cells as presented in Fig. 1B. **(C)** Percentage of γH2aX positive cells from Fig. 1B. Error bars represent the SEM from four independent experiments. Statistical significance was calculated by two-way ANOVA with FDR Benjamini-Hochberg (BH) correction for multiple comparisons (ns = not significant, * = *p* < 0.05). **(D)** Descriptive statistics of microscopy analysis presented in Fig. 1B. **(E)** Total *L. monocytogenes* CFU per spleen extracted from *wildtype* and *Sirt2*^*-/-*^ mice 72 hours post infection.

**Figure S2:**
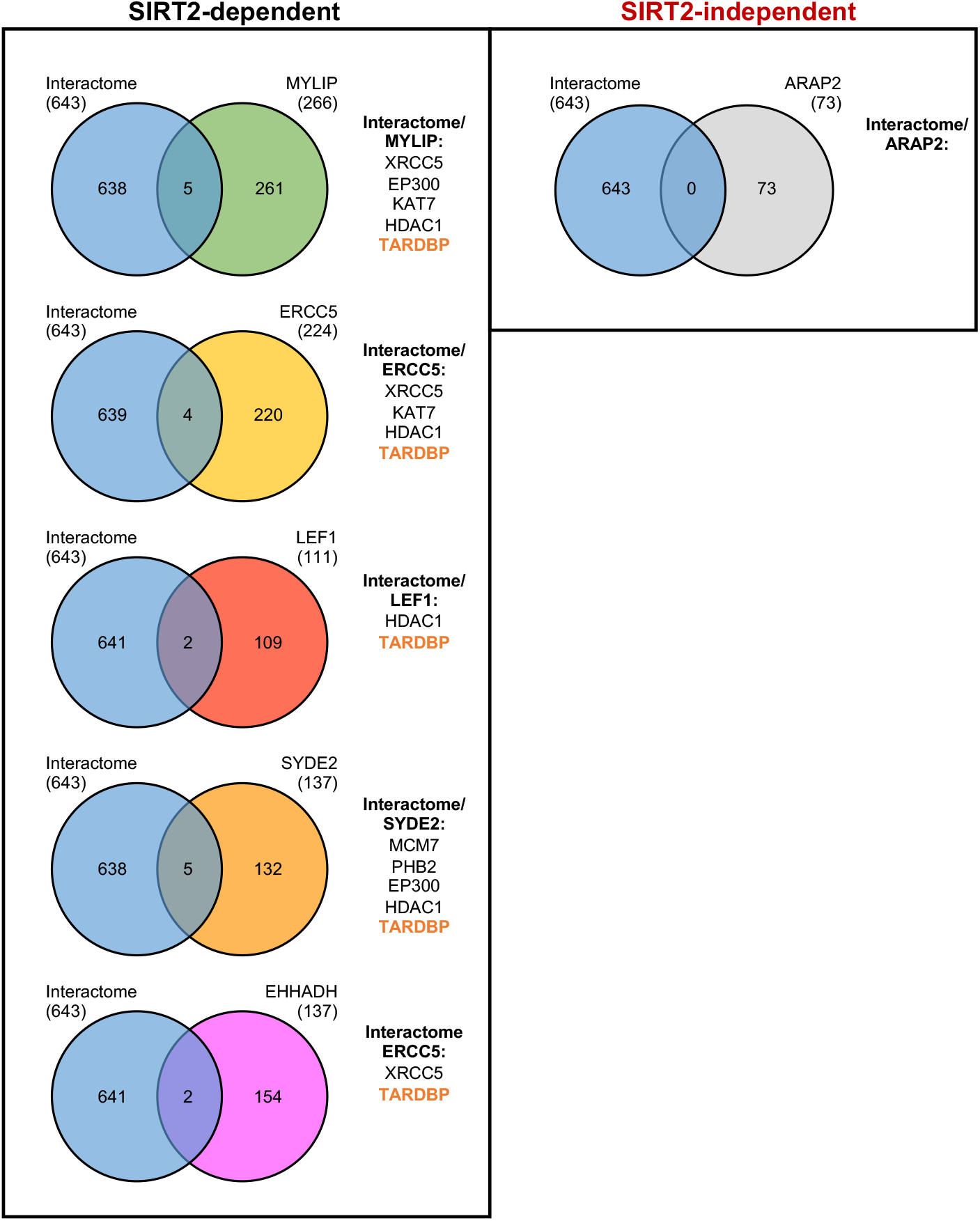
Identification of SIRT2-interacting partners shown to localise to SIRT2-regualted genes by ChIP-seq. Venn diagrams illustrating proteins shared between SIRT2-interactome and interactors of the TSSs of *MYLIP, ERRC5, LEF1, SYDE2, EHHADH* and *ARAP2*.

**Figure S3:**
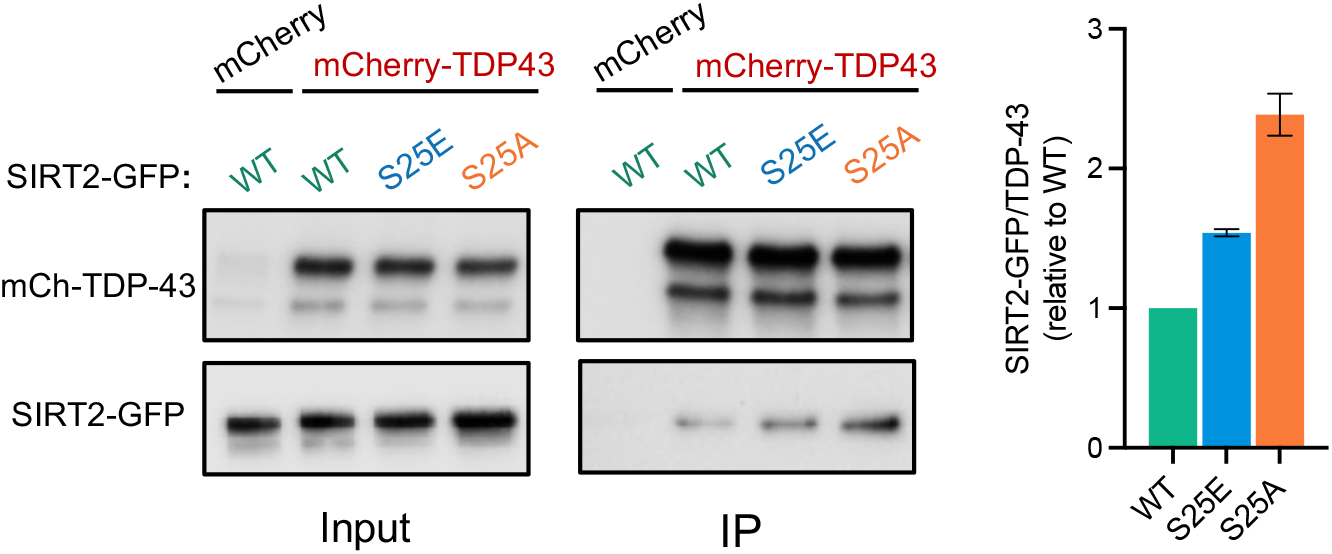
Phosphorylation of SIRT2 at S25 modulates interactions with TDP-43. HeLa cells expressing either mCherry alone or mCherry-TDP-43 were co-transfected with stated variants of SIRT2-GFP followed by immunoprecipitation using RFP-Trap® agarose beads. Cell lysates (Input) and IP fractions were immunoblotted using antibodies against TDP-43 or GFP for detection of SIRT2.

**Figure S4:**
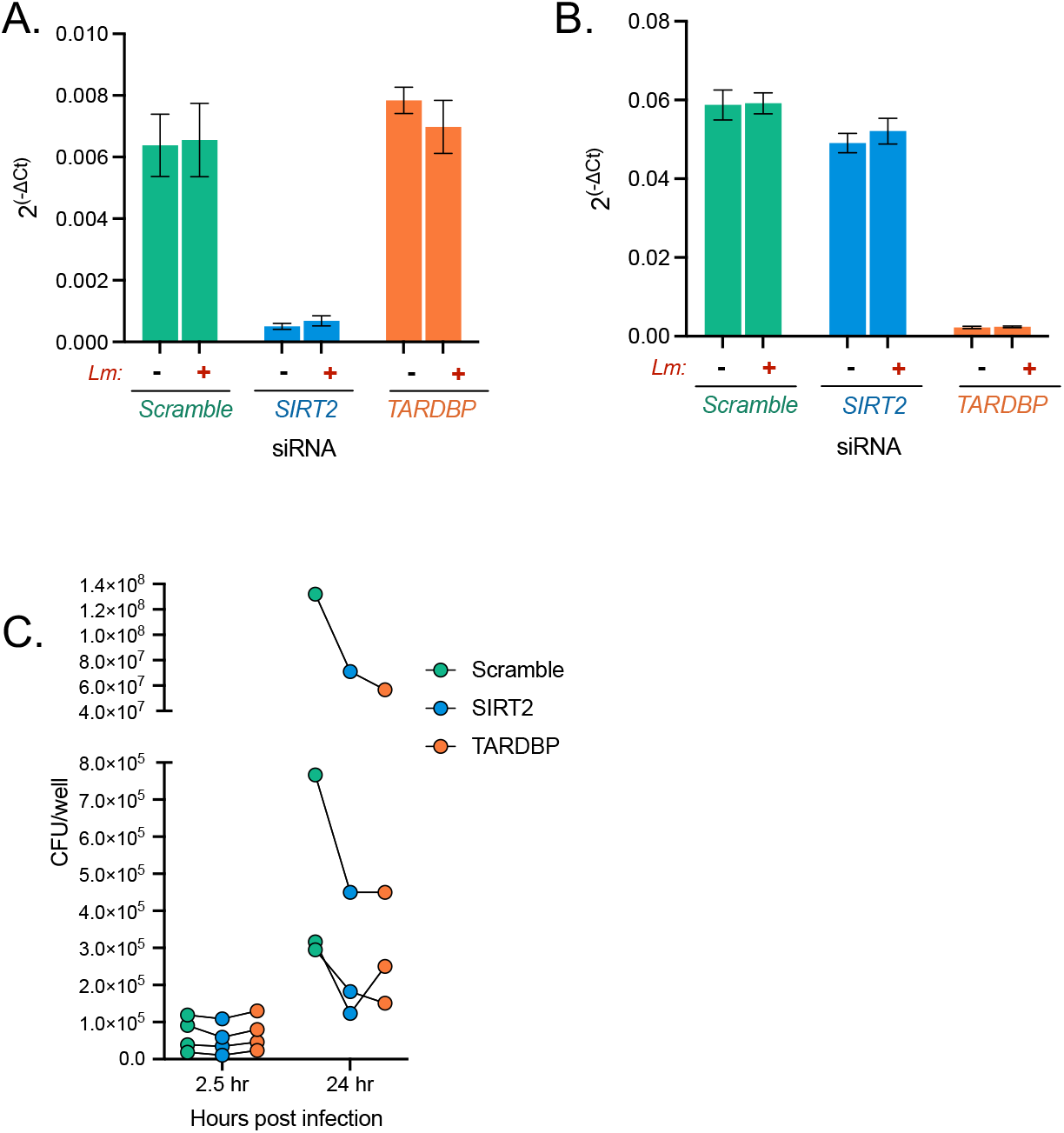
Knockdown of SIRT2 or TDP-43 reduces long term efficacy of *Listeria* infection. Relative mRNA expression of **(A)** *SIRT2* and **(B)** *TARDBP* as detected by qPCR normalised to GAPDH. Mean ± S.E.M from three independent experiments are plotted. **(C)** Quantification of *L. monocytogenes* intracellular CFUs. HeLa cells were transfected with indicated siRNAs and infected for 2.5 h or 24 h. Lysates were plated onto BHI agar and bacterial CFUs were enumerated. Data are presented as CFU/well. Individual biological replicates are plotted as paired values.

**Figure S5:**
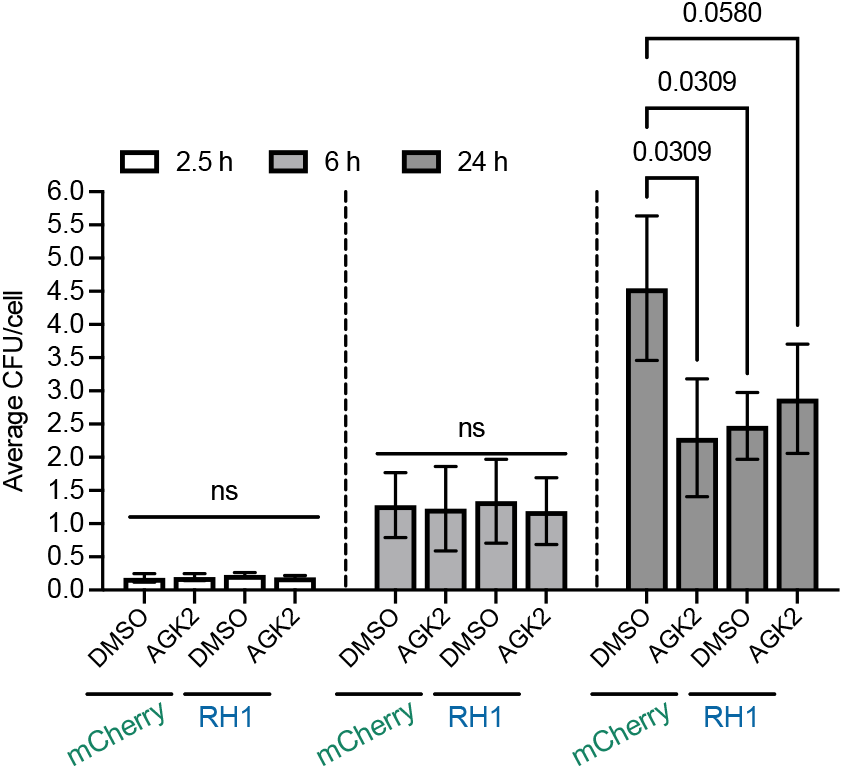
Blocking R-loop formation reduces long term efficacy of *Listeria* infection. Quantification of *L. monocytogenes* intracellular CFU/cell. HeLa cells expressing either mCherry or RNaseH1 were treated with DMSO or 5 mM AGK2 then infected with *L. monocytogenes*. Intracellular bacteria were extracted at 2.5, 6 and 24 hours post infection plated onto BHI agar and bacterial CFUs were enumerated. Data are presented as average CFU/cell. Mean ± S.E.M from three independent experiments are plotted.

## Supplementary Tables

**Table S1:** Bacterial strains used in study

| Bacterial strain | Genotype | Vector | Strain number |
| --- | --- | --- | --- |
| *Listeria monocytogenes* | Wildtype EGD |  | BUG600 |
| *Listeria monocytogenes* | Wildtype EGD | pAD-cGFP | BUG2539 |

**Table S2:** Recipes of buffers used in this study

| Buffer | Components |
| --- | --- |
| Nuclear Isolation buffer | 15 mM Tris (pH 7.5), 60 mM KCl, 15 mM NaCl, 250 mM sucrose, 1 mM CaCl<sub>2</sub>, 5 mM MgCl<sub>2</sub> |
| Chromatin shearing buffer | 1% SDS, 10 mM Tris HCl (pH 8.0), 1 mM EDTA, 0.5 mM EGTA |
| SDS dilution buffer | 0.6% Triton X-100, 0.06% NaDOC, 150 mM NaCl, 12 mM Tris HCl (pH 8.0), 1 mM EDTA, 0.5 mM EGTA |
| Buffer 1 (isotonic) | 1% Triton X-100, 0.1% NaDOC, 150 mM NaCl, 10 mM Tris HCl (pH 8.0); |
| Buffer 2 (isotonic, ionic change) | 0.5% NP40, 0.5% Triton X-100, 0.5% NaDOC, 150 mM NaCl, 10 mM Tris HCl (pH 8.0); |
| Buffer 3 (high salt dilution) | 0.7% Triton X-100, 0.1% NaDOC, 250 mM NaCl, 10 mM Tris HCl (pH 8.0); |
| Buffer 4 (high salt dilution) | 0.5% NP40, 0.5% Triton X-100, 250 mM LiCl, 1 mM EDTA, 20 mM Tris HCl (pH 8.0); |
| Buffer 5 (salt dilution) | 0.1% NP40, 150 mM NaCl, 1 mM EDTA, 20 mM Tris HCl (pH 8.0); |
| Buffer 6 (TE) | 1 mM EDTA, 20 mM Tris HCl (pH 8.0)) |

**Table S3:**
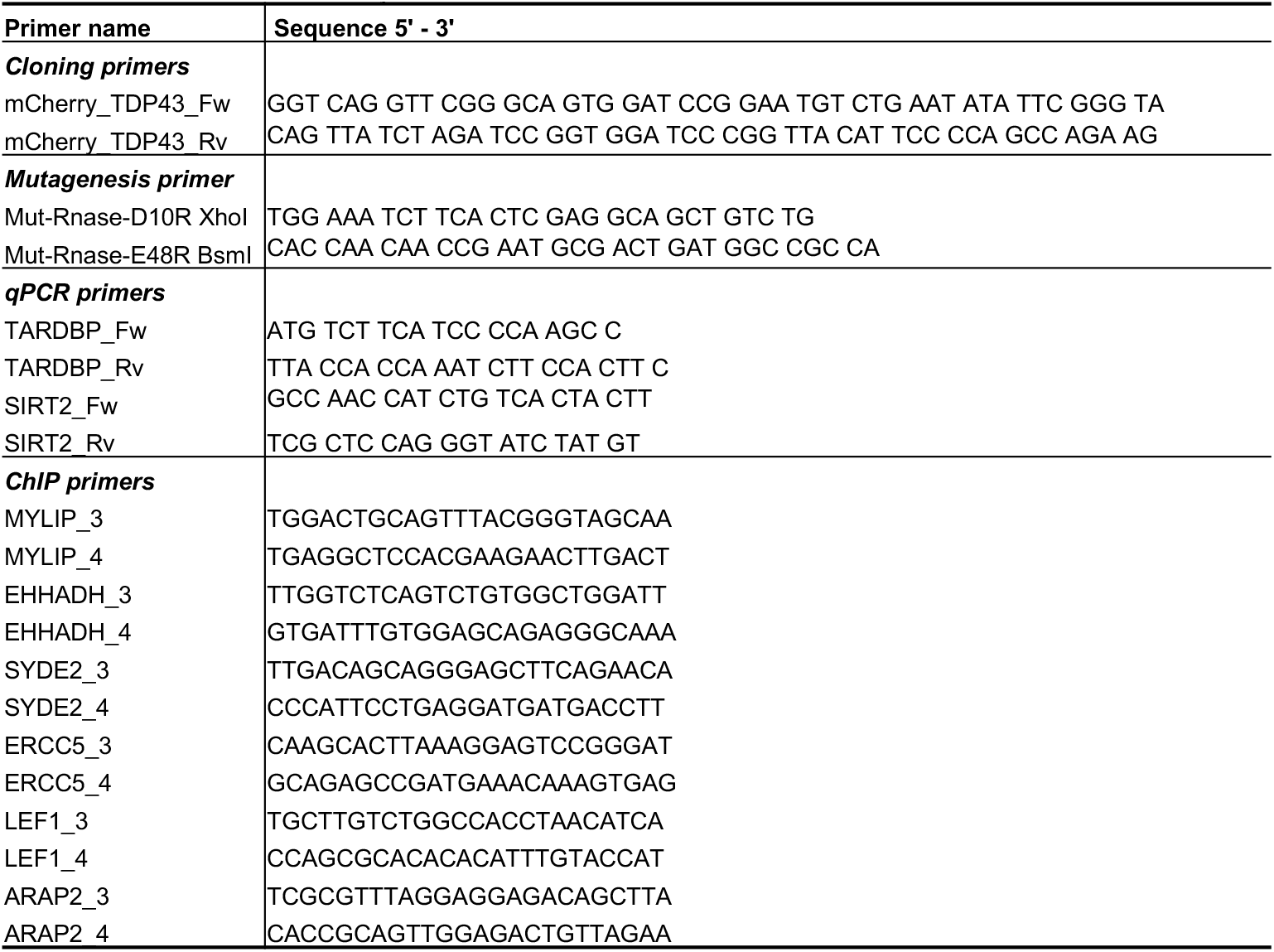
Primers used in this study

**Table S4:** Plasmids used in this study

| Plasmid name | Vector | Origin | Reference |
| --- | --- | --- | --- |
| mCherry-TDP-43 WT | pmCherry-C1 |  | This Study |
| RNaseHI (Dead)-NLS-mCherry | pICE |  | This Study |
| SIRT2-WT-GFP | pEGFP-N1 |  | Pereira et al, 2018 |
| SIRT2-S25A-GFP | pEGFP-N1 |  | Pereira et al, 2018 |
| SIRT2-S25E-GFP | pEGFP-N1 |  | Pereira et al, 2018 |
| H3-GFP WT | pEGFP-N1 | Fang-Lin Sun | Liu et al, 2012 |
| H3-GFP K18A | pEGFP-N1 | Fang-Lin Sun | Liu et al, 2012 |
| H3-GFP K18Q | pEGFP-N1 | Fang-Lin Sun | Liu et al, 2012 |
| NLS-mCherry | pICE | Addgene | Britton et al, 2014 |
| RNaseHI-WT-NLS-mCherry | pICE | Addgene | Britton et al, 2014 |
| TDP43 NOTAG1 | pCAG-EGFP/RFP-int | Addgene | Yang et al, 2010 |

## REFERENCES

Afroz T, Hock E-M, Ernst P, Foglieni C, Jambeau M, Gilhespy LAB, Laferriere F, Maniecka Z, Plückthun A, Mittl P, et al (2017) Functional and dynamic polymerization of the ALS-linked protein TDP-43 antagonizes its pathologic aggregation. Nat Commun 2017 81 8: 1–15

Ashida H, Mimuro H, Ogawa M, Kobayashi T, Sanada T, Kim M & Sasakawa C (2011) Cell death and infection: A double-edged sword for host and pathogen survival. J Cell Biol 195: 931–942

Ayala YM, Zago P, D’Ambrogio A, Xu Y-F, Petrucelli L, Buratti E & Baralle FE (2008) Structural determinants of the cellular localization and shuttling of TDP-43. J Cell Sci 121: 3778–3785

De Bacco F, D’Ambrosio A, Casanova E, Orzan F, Neggia R, Albano R, Verginelli F, Cominelli M, Poliani PL, Luraghi P, et al (2016) MET inhibition overcomes radiation resistance of glioblastoma stem-like cells . EMBO Mol Med 8: 550–568

Behar SM & Briken V (2019) Apoptosis inhibition by intracellular bacteria and its consequence on host immunity. Curr Opin Immunol 60: 103–110

Bhaskar A, Kumar S, Khan MZ, Singh A, Dwivedi VP & Nandicoori VK (2020) Host sirtuin 2 as an immunotherapeutic target against tuberculosis. Elife 9: 1–28

Bierne H & Hamon M (2020) Targeting host epigenetic machinery: The Listeria paradigm. Cell Microbiol 22: e13169

Black JC, Mosley A, Kitada T, Washburn M & Carey M (2008) The SIRT2 Deacetylase Regulates Autoacetylation of p300. Mol Cell 32: 449–455

Bosch-Presegué L & Vaquero A (2014) Sirtuins in stress response: Guardians of the genome. Oncogene 33: 3764–3775

Britton S, Dernoncourt E, Delteil C, Froment C, Schiltz O, Salles B, Frit P & Calsou P (2014) DNA damage triggers SAF-A and RNA biogenesis factors exclusion from chromatin coupled to R-loops removal. Nucleic Acids Res 42: 9047–9062

Buratti E & Baralle FE (2001) Characterization and Functional Implications of the RNA Binding Properties of Nuclear Factor TDP-43, a Novel Splicing Regulator of CFTR Exon 9. J Biol Chem 276: 36337–36343

Buratti E, Brindisi A, Pagani F & Baralle FE (2004) Nuclear factor TDP-43 binds to the polymorphic TG repeats in CFTR intron 8 and causes skipping of exon 9: A functional link with disease penetrance. Am J Hum Genet 74: 1322–1325

Chakraborty A, Tapryal N, Venkova T, Horikoshi N, Pandita RK, Sarker AH, Sarkar PS, Pandita TK & Hazra TK (2016) Classical non-homologous end-joining pathway utilizes nascent RNA for error-free double-strand break repair of transcribed genes. Nat Commun 7: 1–12

Chang C ke, Wu TH, Wu CY, Chiang M hui, Toh EKW, Hsu YC, Lin KF, Liao Y heng, Huang T huang & Huang JJT (2012) The N-terminus of TDP-43 promotes its oligomerization and enhances DNA binding affinity. Biochem Biophys Res Commun 425: 219–224

Chen L, Huang S, Lee L, Davalos A, Schiestl RH, Campisi J & Oshima J (2003) WRN, the protein deficient in Werner syndrome, plays a critical structural role in optimizing DNA repair. Aging Cell 2: 191–199

Cheng Y, Ren X, Gowda ASP, Shan Y, Zhang L, Yuan YS, Patel R, Wu H, Huber-Keener K, Yang JW, et al (2013) Interaction of Sirt3 with OGG1 contributes to repair of mitochondrial DNA and protects from apoptotic cell death under oxidative stress. Cell Death Dis 4: e731–e731

Chumduri C, Gurumurthy RK, Zadora PK, M. Y & Meyer TF (2013) Chlamydia Infection Promotes Host DNA Damage and Proliferation but Impairs the DNA Damage Response. Cell Host Microbe 13: 746–758

Comoglio PM, Trusolino L & Boccaccio C (2018) Known and novel roles of the MET oncogene in cancer: A coherent approach to targeted therapy. Nat Rev Cancer 18: 341–358

Connor MG, Camarasa TMN, Patey E, Rasid O, Barrio L, Weight CM, Miller DP, Heyderman RS, Lamont RJ, Enninga J, et al (2021) The histone demethylase KDM6B fine-tunes the host response to Streptococcus pneumoniae. Nat Microbiol 6: 257–269

Cristini A, Groh M, Kristiansen MS & Gromak N (2018) RNA/DNA Hybrid Interactome Identifies DXH9 as a Molecular Player in Transcriptional Termination and R-Loop-Associated DNA Damage. Cell Rep 23: 1891–1905

Cristini A, Ricci G, Britton S, Salimbeni S, Huang S yin N, Marinello J, Calsou P, Pommier Y, Favre G, Capranico G, et al (2019) Dual Processing of R-Loops and Topoisomerase I Induces Transcription-Dependent DNA Double-Strand Breaks. Cell Rep 28: 3167-3181.e6

Crossley MP, Bocek M & Cimprich KA (2019) R-Loops as Cellular Regulators and Genomic Threats. Mol Cell 73: 398–411

Davis CA, Hitz BC, Sloan CA, Chan ET, Davidson JM, Gabdank I, Hilton JA, Jain K, Baymuradov UK, Narayanan AK, et al (2018) The Encyclopedia of DNA elements (ENCODE): data portal update. Nucleic Acids Res 46: D794–D801

Dong W & Hamon MA (2020) Revealing eukaryotic histone-modifying mechanisms through bacterial infection. Semin Immunopathol 2020 422 42: 201–213

Eldridge MJG, Cossart P & Hamon MA (2020a) Pathogenic Biohacking: Induction, Modulation and Subversion of Host Transcriptional Responses by Listeria monocytogenes. Toxins 2020, Vol 12, Page 294 12: 294

Eldridge MJG, Pereira JM, Impens F & Hamon MA (2020b) Active nuclear import of the deacetylase Sirtuin-2 is controlled by its C-terminus and importins. Sci Rep 10: 1–12

Eskandarian HA, Impens F, Nahori M-A, Soubigou G, Coppée J-Y, Cossart P & Hamon MA (2013) A Role for SIRT2-Dependent Histone H3K18 Deacetylation in Bacterial Infection. Science (80-) 341: 1238858

Fan W & Luo J (2010) SIRT1 regulates UV-induced DNA repair through deacetylating XPA. Mol Cell 39: 247–258

Friedrich A, Pechstein J, Berens C & Lührmann A (2017) Modulation of host cell apoptotic pathways by intracellular pathogens. Curr Opin Microbiol 35: 88–99

Gan W, Guan Z, Liu J, Gui T, Shen K, Manley JL & Li X (2011) R-loop-mediated genomic instability is caused by impairment of replication fork progression. Genes Dev 25: 2041–2056

Gianini M, Bayona-Feliu A, Sproviero D, Barroso SI, Cereda C & Aguilera A (2020) TDP-43 mutations link Amyotrophic Lateral Sclerosis with R-loop homeostasis and R loopmediated DNA damage. PLoS Genet 16: e1009260

Gomes P, Fleming Outeiro T & Cavadas C (2015) Emerging Role of Sirtuin 2 in the Regulation of Mammalian Metabolism. Trends Pharmacol Sci 36: 756–768

Houtkooper RH, Pirinen E & Auwerx J (2012) Sirtuins as regulators of metabolism and healthspan. Nat Rev Mol Cell Biol 13: 225–238

Inoue T, Hiratsuka M, Osaki M, Yamada H, Kishimoto I, Yamaguchi S, Nakano S, Katoh M, Ito H & Oshimura M (2007) SIRT2, a tubulin deacetylase, acts to block the entry to chromosome condensation in response to mitotic stress. Oncogene 26: 945–957

Jeong J-Y, Yim H-S, Ryu J-Y, Lee HS, Lee J-H, Seen D-S & Kang SG (2012) One-step sequence-and ligation-independent cloning as a rapid and versatile cloning method for functional genomics studies. Appl Environ Microbiol 78: 5440–3

Jeong J, Juhn K, Lee H, Kim SH, Min BH, Lee KM, Cho MH, Park GH & Lee KH (2007) SIRT1 promotes DNA repair activity and deacetylation of Ku70. Exp Mol Med 39: 8–13

Jeong SM, Xiao C, Finley LWS, Lahusen T, Souza AL, Pierce K, Li YH, Wang X, Laurent G, German NJ, et al (2013) SIRT4 has tumor-suppressive activity and regulates the cellular metabolic response to dna damage by inhibiting mitochondrial glutamine metabolism. Cancer Cell 23: 450–463

Jo M, Lee S, Jeon YM, Kim S, Kwon Y & Kim HJ (2020) The role of TDP-43 propagation in neurodegenerative diseases: integrating insights from clinical and experimental studies. Exp Mol Med 52: 1652–1662

Kim HS, Patel K, Muldoon-Jacobs K, Bisht KS, Aykin-Burns N, Pennington JD, van der Meer R, Nguyen P, Savage J, Owens KM, et al (2010) SIRT3 Is a Mitochondria-Localized Tumor Suppressor Required for Maintenance of Mitochondrial Integrity and Metabolism during Stress. Cancer Cell 17: 41–52

Kim HS, Vassilopoulos A, Wang RH, Lahusen T, Xiao Z, Xu X, Li C, Veenstra TD, Li B, Yu H, et al (2011) SIRT2 Maintains Genome Integrity and Suppresses Tumorigenesis through Regulating APC/C Activity. Cancer Cell 20: 487–499

Kitamura A, Shibasaki A, Takeda K, Suno R & Kinjo M (2018) Analysis of the substrate recognition state of TDP-43 to single-stranded DNA using fluorescence correlation spectroscopy. Biochem Biophys Reports 14: 58–63

Knodler LA, Finlay B & Steele-Mortimer O (2005) The Salmonella effector protein SopB protects epithelial cells from apoptosis by sustained activation of Akt. J Biol Chem 280: 9058–9064

Konopka A, Whelan DR, Jamali MS, Perri E, Shahheydari H, Toth RP, Parakh S, Robinson T, Cheong A, Mehta P, et al (2020) Impaired NHEJ repair in amyotrophic lateral sclerosis is associated with TDP-43 mutations. Mol Neurodegener 15: 1–28

Kuo PH, Chiang CH, Wang YT, Doudeva LG & Yuan HS (2014) The crystal structure of TDP-43 RRM1-DNA complex reveals the specific recognition for UG-and TG-rich nucleic acids. Nucleic Acids Res 42: 4712–4722

Lagier-Tourenne C, Polymenidou M & Cleveland DW (2010) TDP-43 and FUS/TLS: Emerging roles in RNA processing and neurodegeneration. Hum Mol Genet 19: 46–64

Lalmansingh AS, Urekar CJ & Reddi PP (2011) TDP-43 is a transcriptional repressor: The testis-specific mouse acrv1 gene is a TDP-43 target in vivo. J Biol Chem 286: 10970–10982

Leitão E, Costa AC, Brito C, Costa L, Pombinho R, Cabanes D & Sousa S (2014) Listeria monocytogenes induces host DNA damage and delays the host cell cycle to promote infection. Cell Cycle 13: 928–940

Lemos V, de Oliveira RM, Naia L, Szegö É, Ramos E, Pinho S, Magro F, Cavadas C, Rego AC, Costa V, et al (2017) The NAD+-dependent deacetylase SIRT2 attenuates oxidative stress and mitochondrial dysfunction and improves insulin sensitivity in hepatocytes. Hum Mol Genet 26: 4105–4117

Li HF, Kim JS & Waldman T (2009) Radiation-induced Akt activation modulates radioresistance in human glioblastoma cells. Radiat Oncol 4: 43

Liu Y, Wang DL, Chen S, Zhao L & Sun FL (2012) Oncogene Ras/phosphatidylinositol 3-kinase signaling targets histone H3 acetylation at lysine 56. J Biol Chem 287: 41469–41480

Marnef A & Legube G (2021) R-loops as Janus-faced modulators of DNA repair. Nat Cell Biol 23: 305–313

Medová M, Aebersold DM & Zimmer Y (2014) The molecular crosstalk between the MET receptor tyrosine kinase and the DNA damage response-biological and clinical aspects. Cancers (Basel) 6: 1–27

Mitra J, Guerrero EN, Hegde PM, Liachko NF, Wang H, Vasquez V, Gao J, Pandey A, Paul Taylor J, Kraemer BC, et al (2019) Motor neuron disease-associated loss of nuclear TDP-43 is linked to DNA double-strand break repair defects. Proc Natl Acad Sci U S A 116: 4696–4705

Morales AJ, Carrero JA, Hung PJ, Tubbs AT, Andrews JM, Edelson BT, Calderon B, Innes CL, Paules RS, Payton JE, et al (2017) A type I IFN-dependent DNA damage response regulates the genetic program and inflammasome activation in macrophages. Elife 6

Mosler T, Conte F, Mikicic I, Kreim N, Möckel MM, Flach J, Luke B & Beli P (2021) R-loop proximity proteomics identifies a role of DDX41 in transcription-1 associated genomic instability

Niehrs C & Luke B (2020) Regulatory R-loops as facilitators of gene expression and genome stability. Nat Rev Mol Cell Biol

North BJ & Verdin E (2007) Interphase Nucleo-Cytoplasmic Shuttling and Localization of SIRT2 during Mitosis. PLoS One 2: e784

de Oliveira RM, Sarkander J, Kazantsev AG & Outeiro TF (2012) SIRT2 as a Therapeutic Target for Age-Related Disorders. Front Pharmacol 3: 82

Onn L, Portillo M, Ilic S, Cleitman G, Stein D, Kaluski S, Shirat I, Slobodnik Z, Einav M, Erdel F, et al (2020) SIRT6 is a DNA double-strand break sensor. Elife 9

Peck B, Chen C-Y, Ho K-K, Di Fruscia P, Myatt SS, Coombes RC, Fuchter MJ, Hsiao C-D & Lam EW-F (2010) SIRT inhibitors induce cell death and p53 acetylation through targeting both SIRT1 and SIRT2. Mol Cancer Ther 9: 844–55

Pereira JM, Chevalier C, Chaze T, Gianetto Q, Impens F, Matondo M, Cossart P & Hamon MA (2018) Infection Reveals a Modification of SIRT2 Critical for Chromatin Association. Cell Rep 23: 1124–1137

Pirbhai M, Dong F, Zhong Y, Pan KZ & Zhong G (2006) The Secreted Protease Factor CPAF Is Responsible for Degrading Pro-apoptotic BH3-only Proteins in Chlamydia trachomatis-infected Cells. J Biol Chem 281: 31495–31501

Radoshevich L & Cossart P (2018) Listeria monocytogenes: Towards a complete picture of its physiology and pathogenesis. Nat Rev Microbiol 16: 32–46

Samba-Louaka A, Pereira JM, Nahori MA, Villiers V, Deriano L, Hamon MA & Cossart P (2014) Listeria monocytogenes Dampens the DNA Damage Response. PLoS Pathog 10: 1004470

Serrano L, Martínez-Redondo P, Marazuela-Duque A, Vazquez BN, Dooley SJ, Voigt P, Beck DB, Kane-Goldsmith N, Tong Q, Rabanal RM, et al (2013) The tumor suppressor SirT2 regulates cell cycle progression and genome stability by modulating the mitotic deposition of H4K20 methylation. Genes Dev 27: 639– 653

Shenoy AR & Visweswariah SS (2003) Site-directed mutagenesis using a single mutagenic oligonucleotide and DpnI digestion of template DNA. Anal Biochem 319: 335–336

Tanno M, Sakamoto J, Miura T, Shimamoto K & Horio Y (2007) Nucleocytoplasmic shuttling of the NAD+-dependent histone deacetylase SIRT1. J Biol Chem 282: 6823–32

Tasselli L, Xi Y, Zheng W, Tennen RI, Odrowaz Z, Simeoni F, Li W & Chua KF (2016) SIRT6 deacetylates H3K18ac at pericentric chromatin to prevent mitotic errors and cellular senescence. Nat Struct Mol Biol 23: 434–440

Vaquero A, Scher MB, Dong HL, Sutton A, Cheng HL, Alt FW, Serrano L, Sternglanz R & Reinberg D (2006) SirT2 is a histone deacetylase with preference for histone H4 Lys 16 during mitosis. Genes Dev 20: 1256–1261

Vazquez BN, Thackray JK, Simonet NG, Kane-Goldsmith N, Martinez-Redondo P, Nguyen T, Bunting S, Vaquero A, Tischfield JA & Serrano L (2016) SIRT 7 promotes genome integrity and modulates non-homologous end joining DNA repair . EMBO J 35: 1488–1503

Wang RH, Sengupta K, Li C, Kim HS, Cao L, Xiao C, Kim S, Xu X, Zheng Y, Chilton B, et al (2008) Impaired DNA Damage Response, Genome Instability, and Tumorigenesis in SIRT1 Mutant Mice. Cancer Cell 14: 312–323

Weitzman MD & Weitzman JB (2014) What’s the Damage? The Impact of Pathogens on Pathways that Maintain Host Genome Integrity. Cell Host Microbe 15: 283– 294

Yan F, Cao H, Chaturvedi R, Krishna U, Hobbs SS, Dempsey PJ, Peek RM, Cover TL, Washington MK, Wilson KT, et al (2009) Epidermal Growth Factor Receptor Activation Protects Gastric Epithelial Cells From Helicobacter pylori-Induced Apoptosis. Gastroenterology 136: 1297-1307.e3

Yang C, Tan W, Whittle C, Qiu L, Cao L, Akbarian S & Xu Z (2010) The C-terminal TDP-43 fragments have a high aggregation propensity and harm neurons by a dominant-negative mechanism. PLoS One 5: e15878

Yasuhara T, Kato R, Hagiwara Y, Shiotani B, Yamauchi M, Nakada S, Shibata A & Miyagawa K (2018) Human Rad52 Promotes XPG-Mediated R-loop Processing to Initiate Transcription-Associated Homologous Recombination Repair. Cell 175: 558-570.e11

Zhang H, Park SH, Pantazides BG, Karpiuk O, Warren MD, Hardy CW, Duong DM, Park SJ, Kim HS, Vassilopoulos A, et al (2013) SIRT2 directs the replication stress response through CDK9 deacetylation. Proc Natl Acad Sci U S A 110: 13546–13551

